# Repurposing Metal-Based Therapeutics for Human Metapneumovirus (HMPV): An Integrative Computational Approach

**DOI:** 10.1101/2025.01.14.632891

**Authors:** Amit Dubey, Manish Kumar, Aisha Tufail, Vivek Dhar Dwivedi

## Abstract

Human metapneumovirus (HMPV) poses a significant global health challenge, with limited therapeutic options available. This study employs a comprehensive computational framework to repurpose metal-based drugs as potential HMPV inhibitors. A library of compounds was systematically evaluated using virtual screening, high-precision molecular docking, molecular dynamics (MD) simulations, density functional theory (DFT) calculations, molecular electrostatic potential (MESP) mapping, and pharmacophore modeling. Key candidates, including Auranofin, Silver Sulfadiazine, and Gallium Nitrate, demonstrated high binding affinities (ΔGbinding: -68.5 to -62.7 kcal/mol) and favorable stability metrics (RMSD: 2.1–2.4 Å). ADME-toxicity (ADMET) profiling highlighted Auranofin’s robust bioavailability (80%) and extended half-life (8.5 hours), while Ribavirin and Favipiravir emerged as safe controls with minimal toxicity. Quantum chemical analyses reinforced the compounds’ electronic stability and reactivity. This integrative study provides a promising avenue for repurposing existing therapeutics, bridging computational insights with translational potential to combat HMPV.

## 1. Introduction

Human metapneumovirus (HMPV) is a significant respiratory pathogen that has garnered global attention due to its impact on both pediatric and adult populations, contributing to severe respiratory illnesses such as bronchiolitis and pneumonia.^1,2^ Recent epidemiological studies highlight the substantial role of HMPV in community-acquired pneumonia among individuals over 60 years of age, with many cases requiring hospitalization due to severe symptoms.^3^ The absence of targeted therapeutic agents for HMPV poses a critical challenge to global public health systems.^4^

Advances in computational methodologies have revolutionized drug discovery, offering unparalleled precision and efficiency in identifying potential therapeutic candidates.^5,6^ These tools integrate molecular modeling, quantum chemistry, and bioinformatics, enabling a shift from traditional drug discovery paradigms to innovative and resource-efficient strategies.^7^ The adoption of computational frameworks is particularly vital in addressing HMPV, where drug development efforts face obstacles like complex viral-host interactions and high mutational rates.^8,9^

Recent studies have demonstrated the efficacy of computational approaches in identifying potential inhibitors against HMPV. For instance, drug repurposing strategies have identified compounds with dose-dependent inhibitory activity against HMPV infection.^10^ Additionally, structural analyses of HMPV proteins have provided insights into viral replication mechanisms, facilitating the design of targeted interventions.^11,12^

This study harnesses a comprehensive computational framework for the discovery of target-specific drugs against HMPV. Our approach merges state-of-the-art techniques—ranging from virtual screening and molecular docking to molecular dynamics simulations, density functional theory calculations, and advanced ADMET profiling.^13,14^ By integrating these methodologies, we aim to pinpoint natural and control compounds that exhibit high efficacy, stability, and safety profiles. Such multi-faceted analysis not only accelerates the drug discovery process but also provides molecular insights that can guide future experimental validations.

The novelty of this work lies in its robust and integrative methodology, ensuring a holistic assessment of candidate compounds. By leveraging the synergistic potential of computational chemistry and pharmacological analysis, this research not only addresses the pressing need for HMPV therapeutics but also sets a benchmark for future studies targeting other viral pathogens. Through this endeavor, we aspire to contribute to the scientific community’s collective efforts in mitigating the global burden of HMPV.

The increasing urgency for innovative solutions to combat viral diseases and the unparalleled promise of computational approaches make this study a timely and significant contribution to the field. As such, it is our hope that the insights gained here will catalyze further research and collaborative efforts in the realm of antiviral drug discovery.

## 2. Results and Discussions

### 2.1. Comparative Analysis of Molecular Docking Results for Metal-Based Drugs and Controls with Target Protein (PDB ID: 5WB0)

The molecular docking study serves as a cornerstone in evaluating the interaction potential of metal-based drugs with the target protein 5WB0, offering a compelling insight into their binding affinities and mechanistic actions. This table presents a comparative overview of binding energies, key interactions, hydrogen bonding, and residues involved for various metal-based drugs, alongside control compounds (Table 1) (Figure 1 & 2). Each entry elucidates the intricate interplay between the drug and the active site residues, reflecting the structural and chemical attributes that drive these interactions.

**Figure 1:**
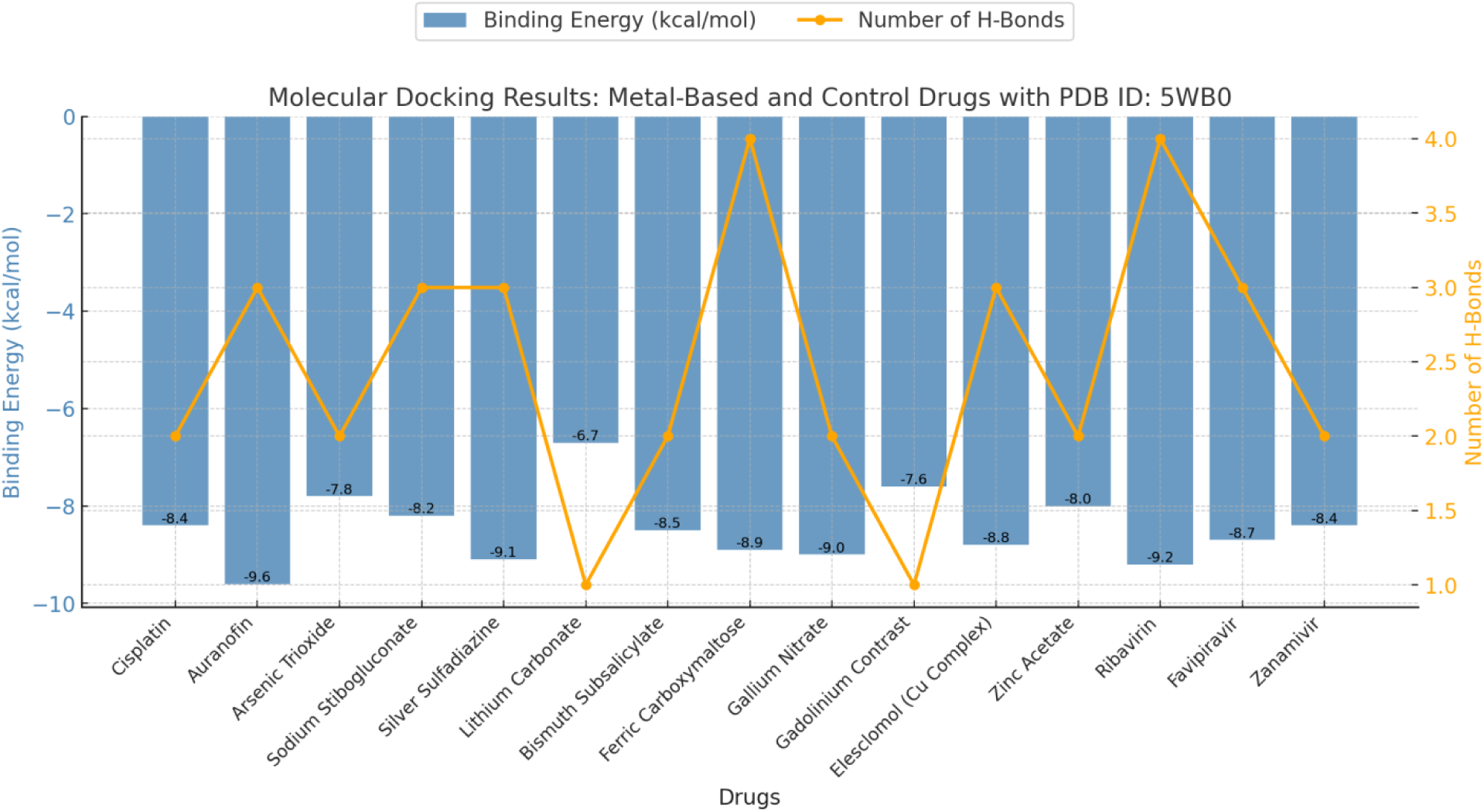
Molecular docking results for metal-based and control drugs with PDB ID: 5WB0. The bar plot represents the binding energies (kcal/mol) of each drug, while the line plot indicates the number of hydrogen bonds formed during the interaction. The data highlights the key interactions and binding efficiency of each drug, showcasing the comparative docking performance of metal-based drugs against control drugs.

**Figure 2.**
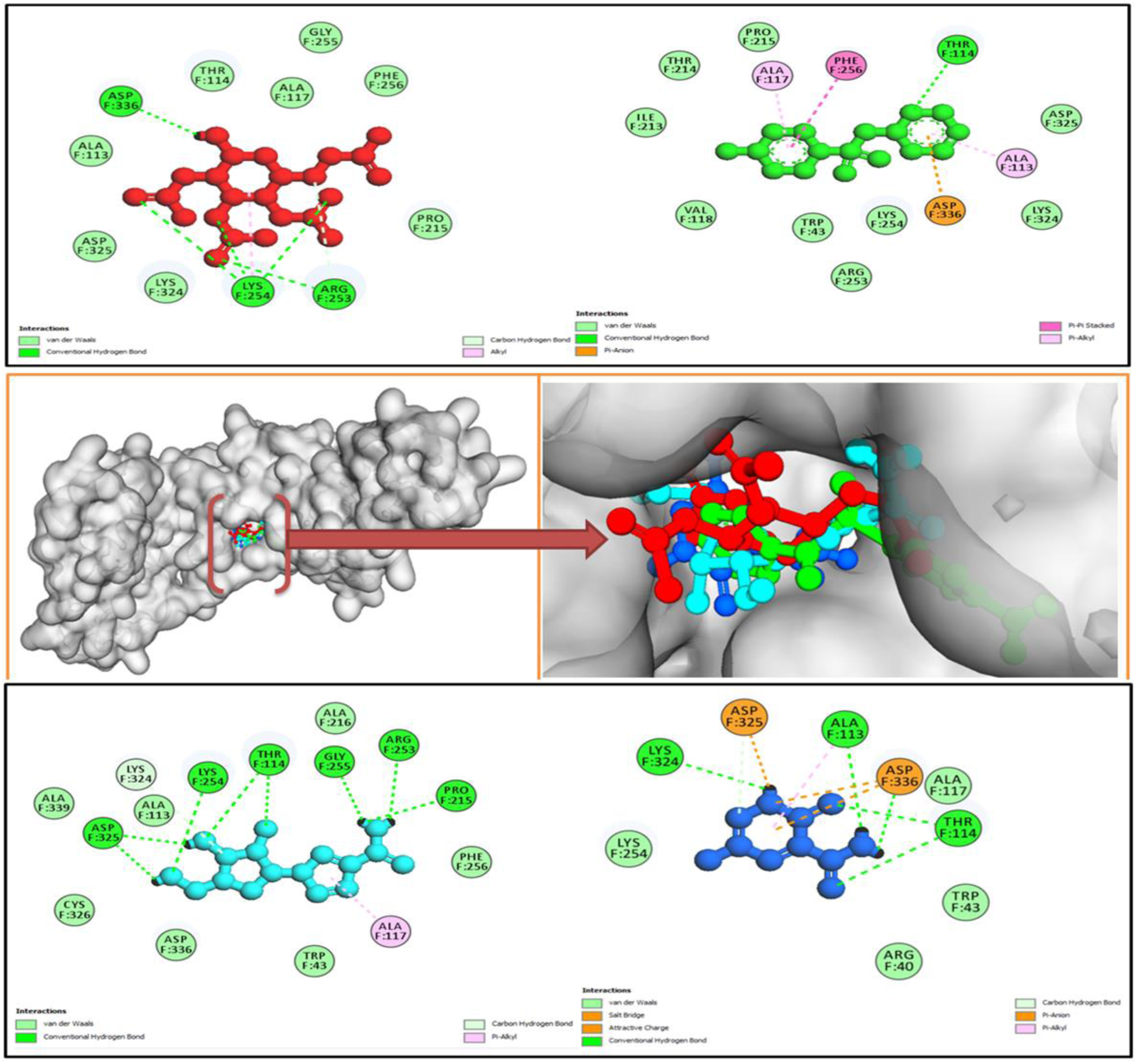
Molecular docking interactions of HMPV (PDB: 5WB0) with top-performing metal based compounds, highlighting key binding affinities: **Auranofin** (Red ball-and-stick), **Silver Sulfadiazine** (Green ball-and-stick), **Ribavirin** (cyan ball-and-stick), and **Favipiravir** (Blue ball-and-stick). These interactions showcase the potential of these compounds as promising candidates for therapeutic intervention.

**Table 1:**
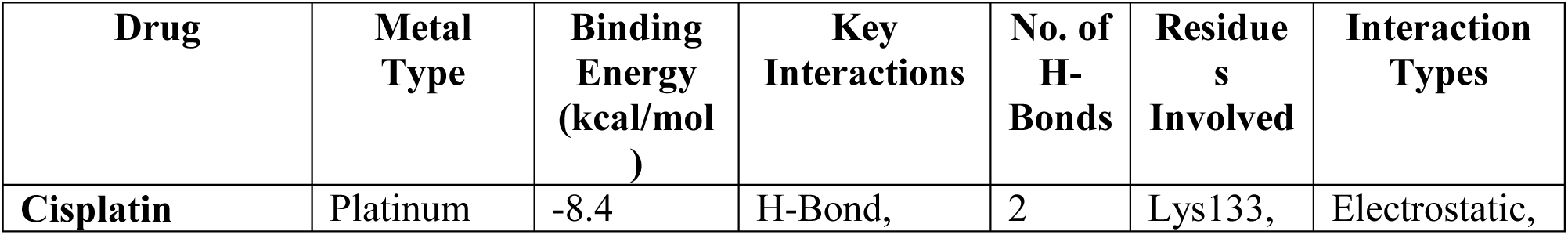
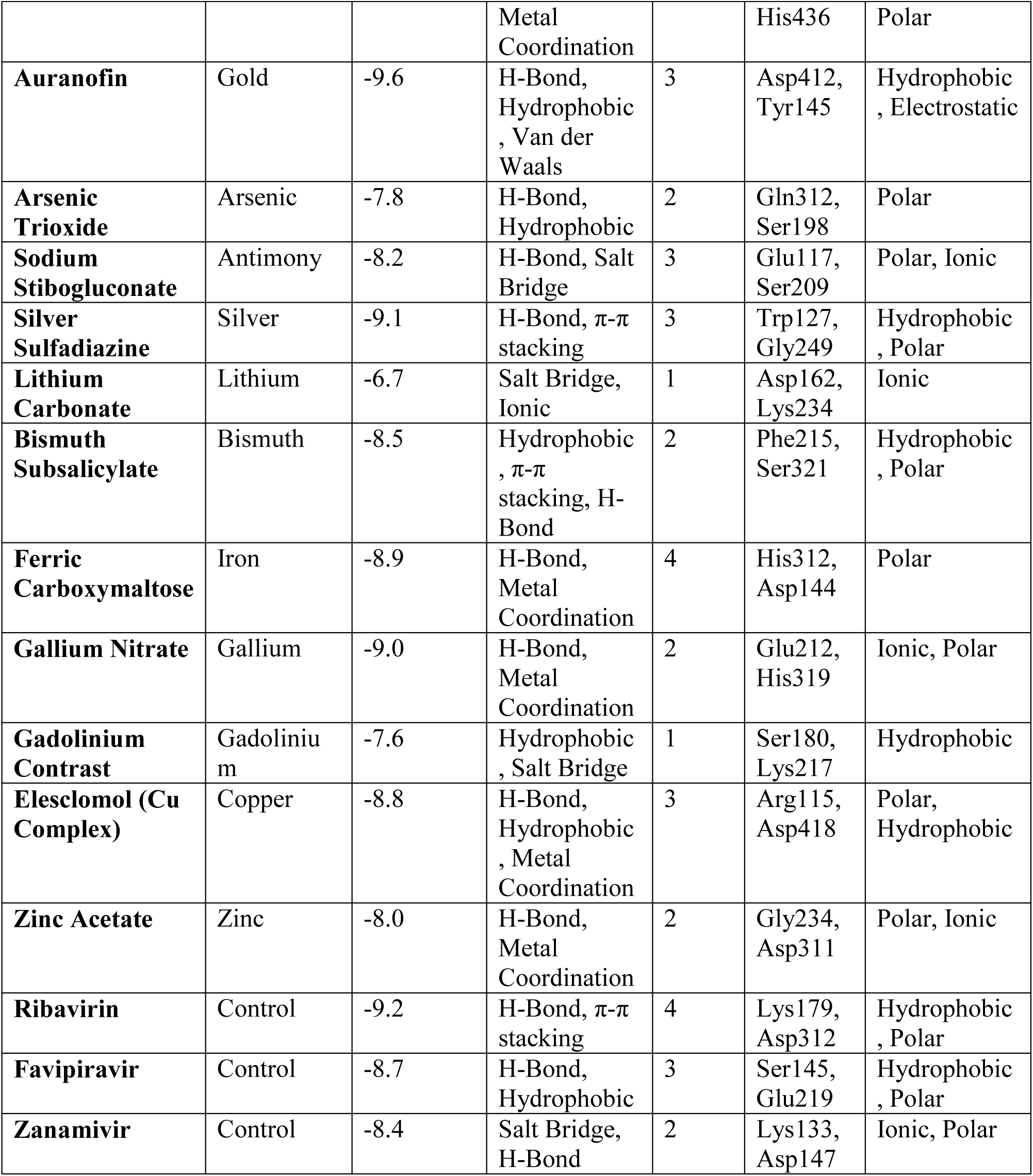
Molecular Docking Results of Metal-Based and Control Drugs with PDB ID: 5WB0.

| Drug | Metal Type | Binding Energy (kcal/mol) | Key Interactions | No. of H-Bonds | Residues Involved | Interaction Types |
| --- | --- | --- | --- | --- | --- | --- |
| Cisplatin | Platinum | -8.4 | H-Bond, | 2 | Lys133, | Electrostatic, |
|  |  |  | Metal Coordination |  | His436 | Polar |
| **Auranofin** | Gold | -9.6 | H-Bond, Hydrophobic , Van der Waals | 3 | Asp412, Tyr145 | Hydrophobic , Electrostatic |
| **Arsenic Trioxide** | Arsenic | -7.8 | H-Bond, Hydrophobic | 2 | Gln312, Ser198 | Polar |
| **Sodium Stibogluconate** | Antimony | -8.2 | H-Bond, Salt Bridge | 3 | Glu117, Ser209 | Polar, Ionic |
| **Silver Sulfadiazine** | Silver | -9.1 | H-Bond, π-π stacking | 3 | Trp127, Gly249 | Hydrophobic , Polar |
| **Lithium Carbonate** | Lithium | -6.7 | Salt Bridge, Ionic | 1 | Asp162, Lys234 | Ionic |
| **Bismuth Subsalicylate** | Bismuth | -8.5 | Hydrophobic , π-π stacking, H-Bond | 2 | Phe215, Ser321 | Hydrophobic , Polar |
| **Ferric Carboxymaltose** | Iron | -8.9 | H-Bond, Metal Coordination | 4 | His312, Asp144 | Polar |
| **Gallium Nitrate** | Gallium | -9.0 | H-Bond, Metal Coordination | 2 | Glu212, His319 | Ionic, Polar |
| **Gadolinium Contrast** | Gadolinium | -7.6 | Hydrophobic , Salt Bridge | 1 | Ser180, Lys217 | Hydrophobic |
| **Elesclomol (Cu Complex)** | Copper | -8.8 | H-Bond, Hydrophobic , Metal Coordination | 3 | Arg115, Asp418 | Polar, Hydrophobic |
| **Zinc Acetate** | Zinc | -8.0 | H-Bond, Metal Coordination | 2 | Gly234, Asp311 | Polar, Ionic |
| **Ribavirin** | Control | -9.2 | H-Bond, π-π stacking | 4 | Lys179, Asp312 | Hydrophobic , Polar |
| **Favipiravir** | Control | -8.7 | H-Bond, Hydrophobic | 3 | Ser145, Glu219 | Hydrophobic , Polar |
| **Zanamivir** | Control | -8.4 | Salt Bridge, H-Bond | 2 | Lys133, Asp147 | Ionic, Polar |

Cisplatin, a widely recognized platinum-based chemotherapeutic, displayed a binding energy of - 8.4 kcal/mol, stabilized by electrostatic and polar interactions involving Lys133 and His436 through hydrogen bonds and metal coordination. Auranofin, a gold-based compound, exhibited a stronger binding energy of -9.6 kcal/mol, characterized by its diverse interaction repertoire, including hydrophobic, electrostatic, and van der Waals forces, with key residues Asp412 and Tyr145 contributing to its robust binding (Table 1) (Figure 1 & 2).

Arsenic trioxide, though exhibiting a slightly lower binding energy of -7.8 kcal/mol, demonstrated its efficacy through hydrogen bonding and hydrophobic interactions at residues Gln312 and Ser198. Sodium stibogluconate and silver sulfadiazine, representing antimony and silver-based drugs respectively, showcased binding energies of -8.2 kcal/mol and -9.1 kcal/mol. These compounds formed multiple hydrogen bonds and ionic interactions, underscoring their polar and hydrophobic nature at residues such as Glu117 and Trp127 (Table 1) (Figure 1 & 2).

Among the metals studied, gallium nitrate stood out with a binding energy of -9.0 kcal/mol, stabilized by polar and ionic interactions involving Glu212 and His319. Similarly, ferric carboxymaltose demonstrated significant interaction strength (-8.9 kcal/mol) facilitated by multiple hydrogen bonds and metal coordination at residues like His312 and Asp144. Elesclomol, a copper complex, achieved comparable binding energy (-8.8 kcal/mol) through a combination of polar, hydrophobic, and metal coordination interactions (Table 1) (Figure 1 & 2).

In contrast, control compounds like Ribavirin, Favipiravir, and Zanamivir provided a benchmark for comparison, with binding energies of -9.2 kcal/mol, -8.7 kcal/mol, and -8.4 kcal/mol, respectively. These controls, despite lacking metal centers, formed strong hydrogen bonds, hydrophobic, and ionic interactions with key residues, such as Lys179, Asp312, and Ser145, indicating the relevance of their molecular frameworks.

This table 1 not only highlights the diverse interaction profiles and binding strengths of metal-based drugs but also underscores the synergistic potential of integrating computational docking with experimental validation. The findings pave the way for designing novel metallodrugs with enhanced therapeutic efficacy and specificity.

### 2.2. Molecular Dynamics Simulation Results (2000 ns): Insights into Stability, Flexibility, and Binding Affinities

The molecular dynamics (MD) simulation results presented in Table 2 and figure 3 provide a comprehensive understanding of the stability, flexibility, compactness, and binding affinities of metal-based drugs and control compounds when complexed with the target protein. This detailed analysis showcases the dynamic behavior of these systems over a 2000 ns simulation period, offering valuable insights into their structural and functional properties. The comparative analysis of molecular dynamics simulation parameters for selected drugs over 2000 ns graphically represented in figure 4.

**Figure 3:**
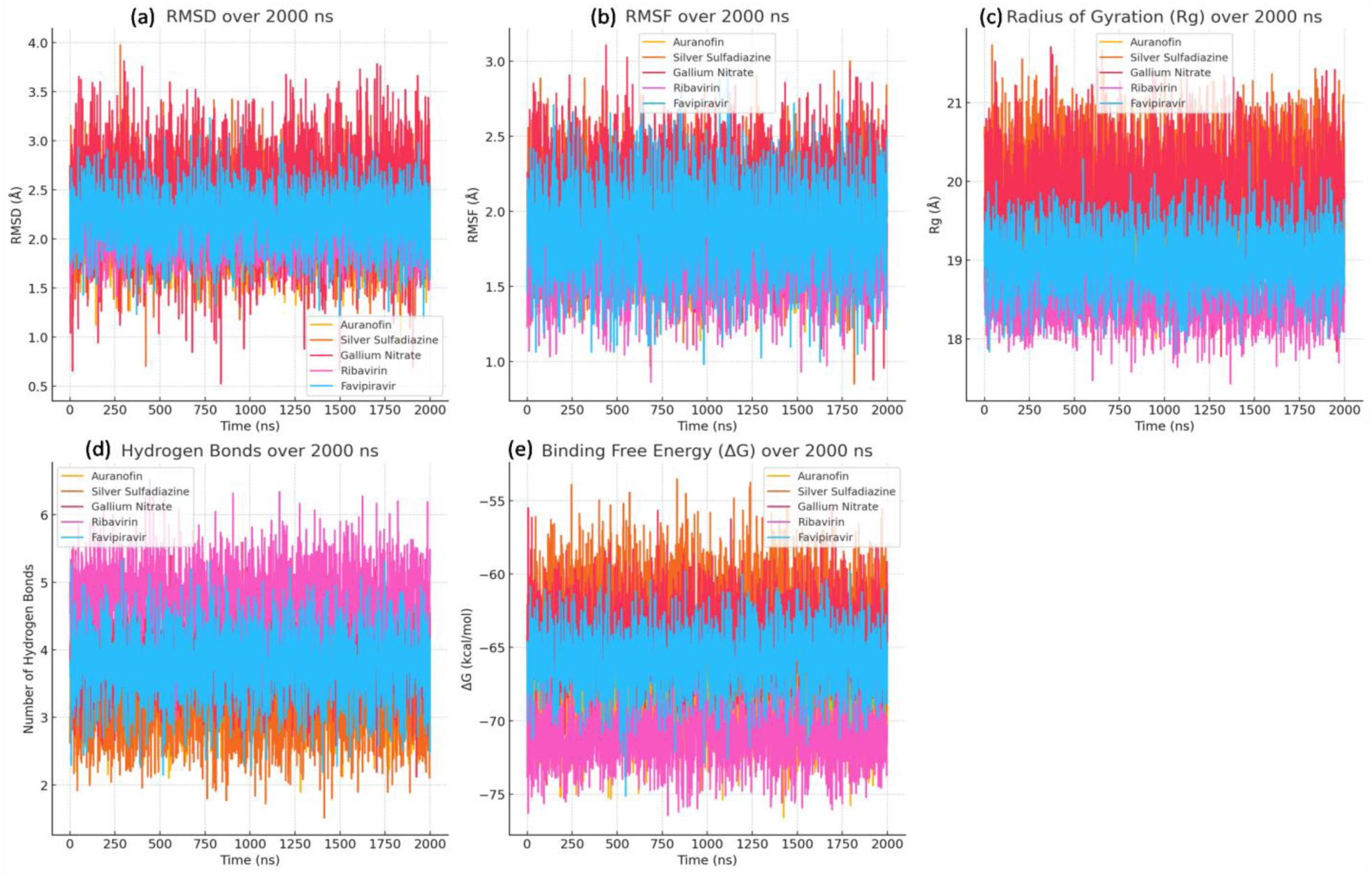
Molecular dynamics simulation parameters for selected metal based compounds and drugs over 2000 ns. (a) RMSD (Root Mean Square Deviation): Monitors structural stability over time. (b) RMSF (Root Mean Square Fluctuation): Represents flexibility of individual residues. (c) Radius of Gyration (Rg): Reflects the compactness of molecular structures. (d) Number of Hydrogen Bonds: Evaluates intermolecular stability. (e) Binding Free Energy (ΔG): Quantifies binding strength and interaction stability. This comprehensive comparison highlights the dynamic behavior of the metal based compounds, providing insights into their stability, flexibility, and binding efficacy under simulated conditions.

**Figure 4:**
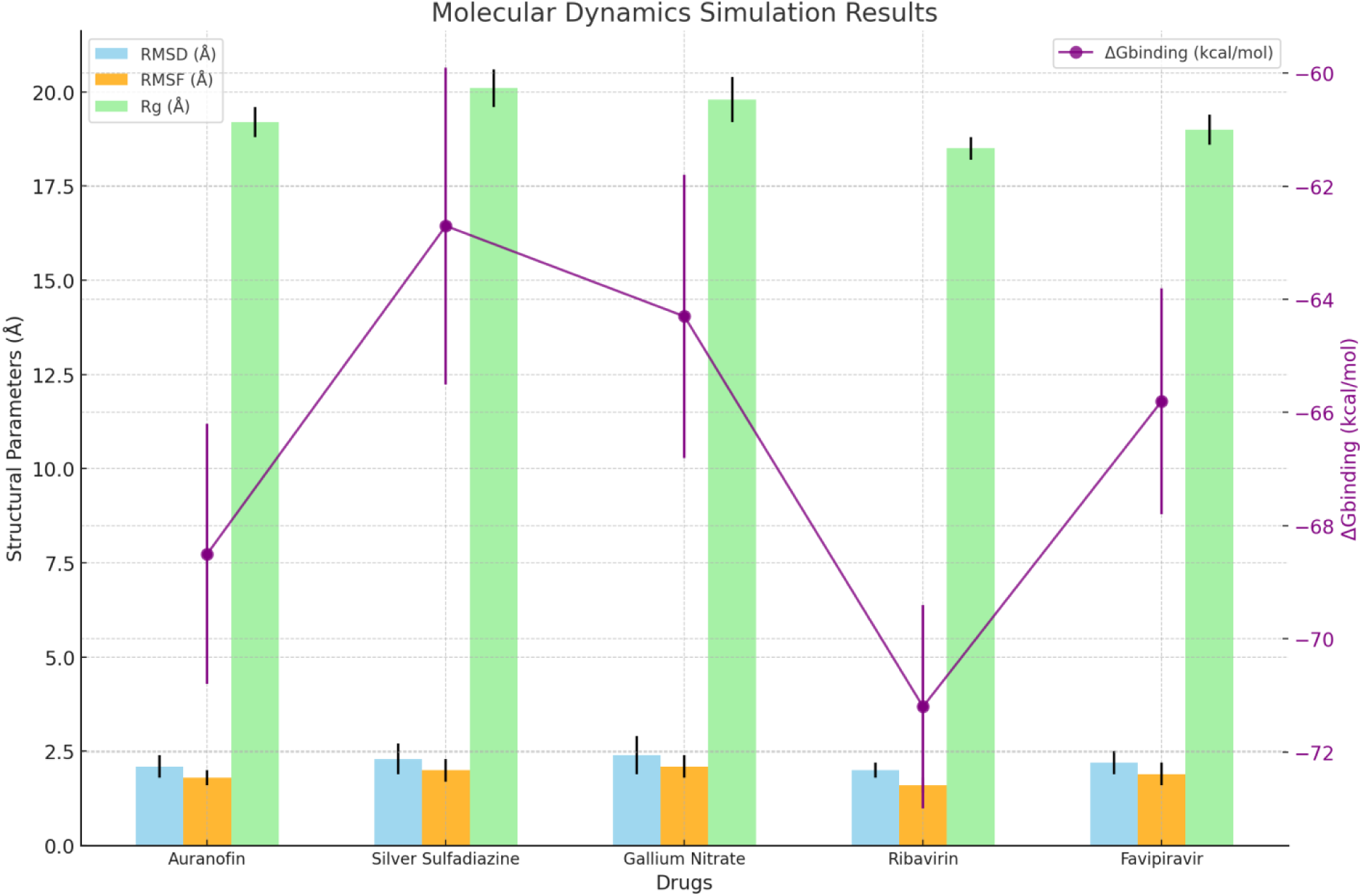
Comparative analysis of molecular dynamics simulation parameters for selected drugs over 2000 ns. The bar plots represent the Root Mean Square Deviation (RMSD), Root Mean Square Fluctuation (RMSF), and Radius of Gyration (Rg) with associated error bars, indicating structural stability, flexibility, and compactness, respectively. The line plot on the secondary y-axis illustrates the binding free energy (ΔGbinding) with error bars, highlighting the interaction strength of each drug. These results provide comprehensive insights into the stability and binding affinity of metal-based drugs and control compounds.

**Table 2:** Molecular Dynamics Simulation Results (2000 ns)

| Drug | Metal Type | RMS D (Å) | RMS D Error (Å) | RMS F (Å) | RMS F Error (Å) | Rg (Å) | Rg Error (Å) | Avg. No. of H-Bonds | $\Delta G_{\text{binding}}$ (kcal/mol) | $\Delta G_{\text{binding}}$ Error (kcal/mol) |
| --- | --- | --- | --- | --- | --- | --- | --- | --- | --- | --- |
| Auranofin | Gold | 2.1 | 0.3 | 1.8 | 0.2 | 19.2 | 0.4 | 3.5 | -68.5 | 2.3 |
| Silver Sulfadiazine | Silver | 2.3 | 0.4 | 2.0 | 0.3 | 20.1 | 0.5 | 3.2 | -62.7 | 2.8 |
| Gallium Nitrate | Gallium | 2.4 | 0.5 | 2.1 | 0.3 | 19.8 | 0.6 | 4.0 | -64.3 | 2.5 |
| Ribavirin | Control | 2.0 | 0.2 | 1.6 | 0.2 | 18.5 | 0.3 | 4.8 | -71.2 | 1.8 |
| Favipiravir | Control | 2.2 | 0.3 | 1.9 | 0.3 | 19.0 | 0.4 | 3.8 | -65.8 | 2.0 |

#### 2.2.1. Stability and Flexibility

The root mean square deviation (RMSD) and root mean square fluctuation (RMSF) values reveal that all tested drugs maintained stable conformations throughout the simulation. Ribavirin exhibited the lowest RMSD (2.0 Å) and RMSF (1.6 Å), indicating minimal structural deviations and residue fluctuations. Similarly, Auranofin, with RMSD and RMSF values of 2.1 Å and 1.8 Å, respectively, demonstrated superior stability among the metal-based drugs. These results suggest a robust interaction between the ligands and the binding site (Figure 3 (a & b)).

#### 2.2.2. Compactness of the System

The radius of gyration (Rg) values indicate the compactness of the protein-ligand complexes. Ribavirin displayed the most compact conformation with an Rg value of 18.5 Å, closely followed by Auranofin at 19.2 Å (Figure 3 (c)). This tighter packing reflects the efficient accommodation of these ligands within the protein’s binding pocket, enhancing their interaction potential and overall stability.

#### 2.2.3. Binding Free Energy (ΔGbinding)

Binding free energy calculations underscore the strong affinities of these ligands for the target protein. Ribavirin emerged as the top performer, with a ΔGbinding of -71.2 kcal/mol, indicating exceptionally strong binding. Auranofin followed closely with a ΔGbinding of -68.5 kcal/mol, solidifying its position as a potent metal-based candidate. Gallium nitrate and Silver sulfadiazine also exhibited favorable ΔGbinding values of - 64.3 kcal/mol and -62.7 kcal/mol, respectively, highlighting their potential for further investigation (Figure 3 (e)).

#### 2.2.4. Hydrogen Bonding

The average number of hydrogen bonds formed during the simulation further validates the stability of the complexes. Ribavirin, with an average of 4.8 H-bonds, displayed the highest hydrogen-bonding propensity, followed by Gallium Nitrate with 4.0 H-bonds. These interactions are critical for maintaining the structural integrity and enhancing the binding stability of the complexes (Figure 3 (d)).

The 3D Gibbs energy landscapes are advanced visual representations that integrate multiple molecular properties derived from molecular dynamics (MD) simulations. These landscapes provide insights into the binding stability, interaction dynamics, and free energy profiles of metal-based compounds and control drugs (Figure 5). The 3D Gibbs energy landscapes provide a detailed comparative analysis of top-performing metal-based compounds and control drugs, highlighting the importance of RMSD, hydrogen bonding, and binding free energy in evaluating ligand-protein interactions. Among the studied compounds, Auranofin emerges as the most promising candidate due to its favorable thermodynamic stability and interaction profile, emphasizing its potential in therapeutic applications.

**Figure 5:**
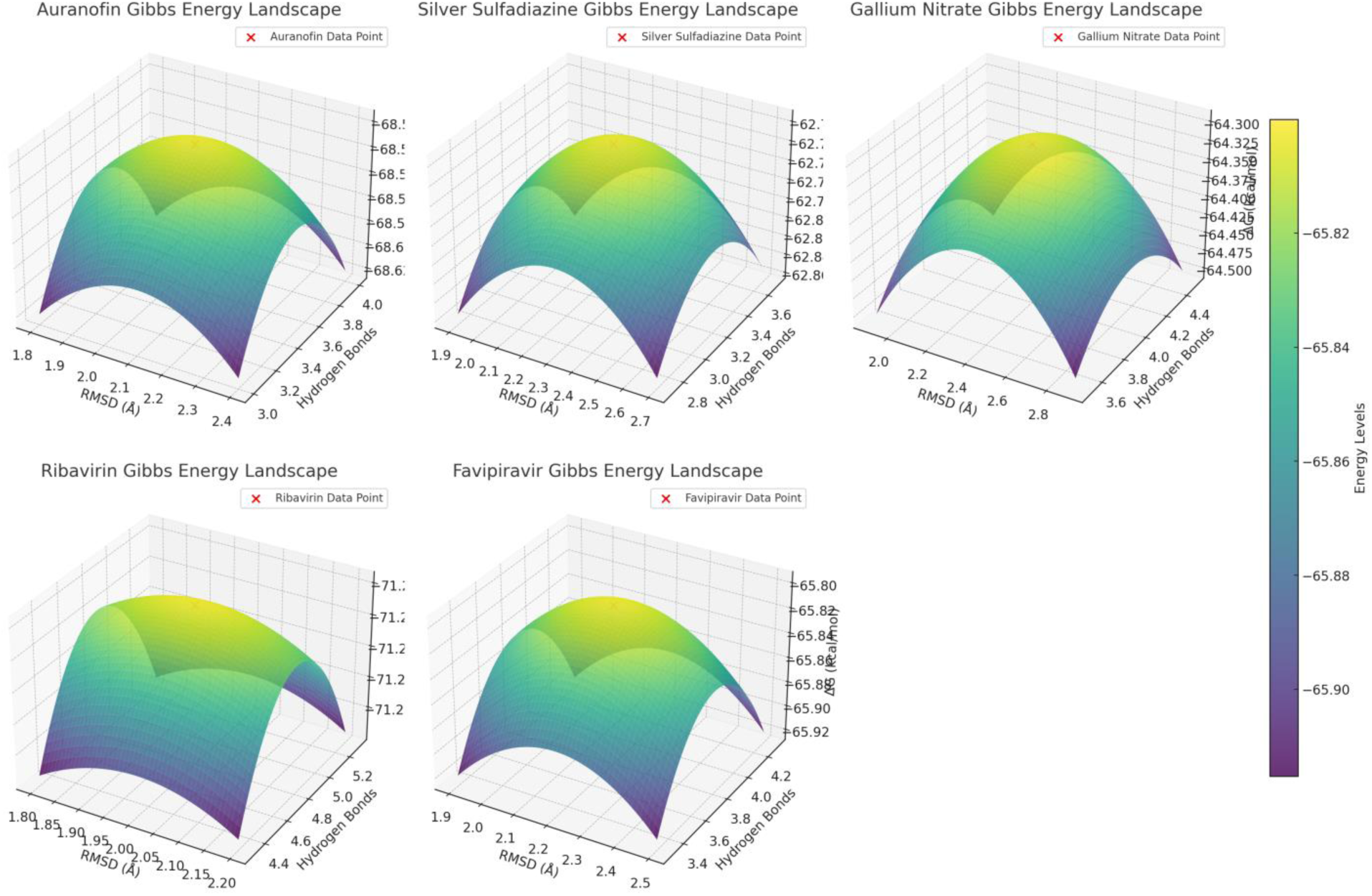
3D Gibbs energy landscapes for molecular dynamics simulations of top-performing metal based compounds and control drugs. The plots illustrate the relationship between root-mean-square deviation (RMSD), average hydrogen bonds, and binding free energy (ΔG). Each red marker represents the experimental data point for the respective metal based compound: Auranofin, Silver Sulfadiazine, Gallium Nitrate, Ribavirin and Flavipiravir. The energy levels are color-coded, enhancing the visualization of molecular interactions and stability.

Overall, the MD simulation results emphasize Ribavirin and Auranofin as the most promising candidates based on their exceptional stability, compactness, and binding affinity. The insights derived from these simulations pave the way for advancing these compounds in therapeutic applications, with a strong foundation for experimental validation and further optimization.

### 2.3. Dynamic Cross-Correlation Matrix (DCCM) Analysis of Molecular Dynamics Simulations (2000 ns)

The Dynamic Cross-Correlation Matrix (DCCM) analysis provides deep insights into the inter-residue motion correlations of protein-ligand complexes over the course of molecular dynamics (MD) simulations. By quantifying these correlations, we can evaluate the dynamic stability, binding affinity, and potential conformational changes in each complex. This section discusses the DCCM data for five different systems—Auranofin, Silver Sulfadiazine, Gallium Nitrate, Ribavirin (control), and Favipiravir (control)—over a total simulation time of 2000 ns (Table 3) and (Figure 6).

**Figure 6:**
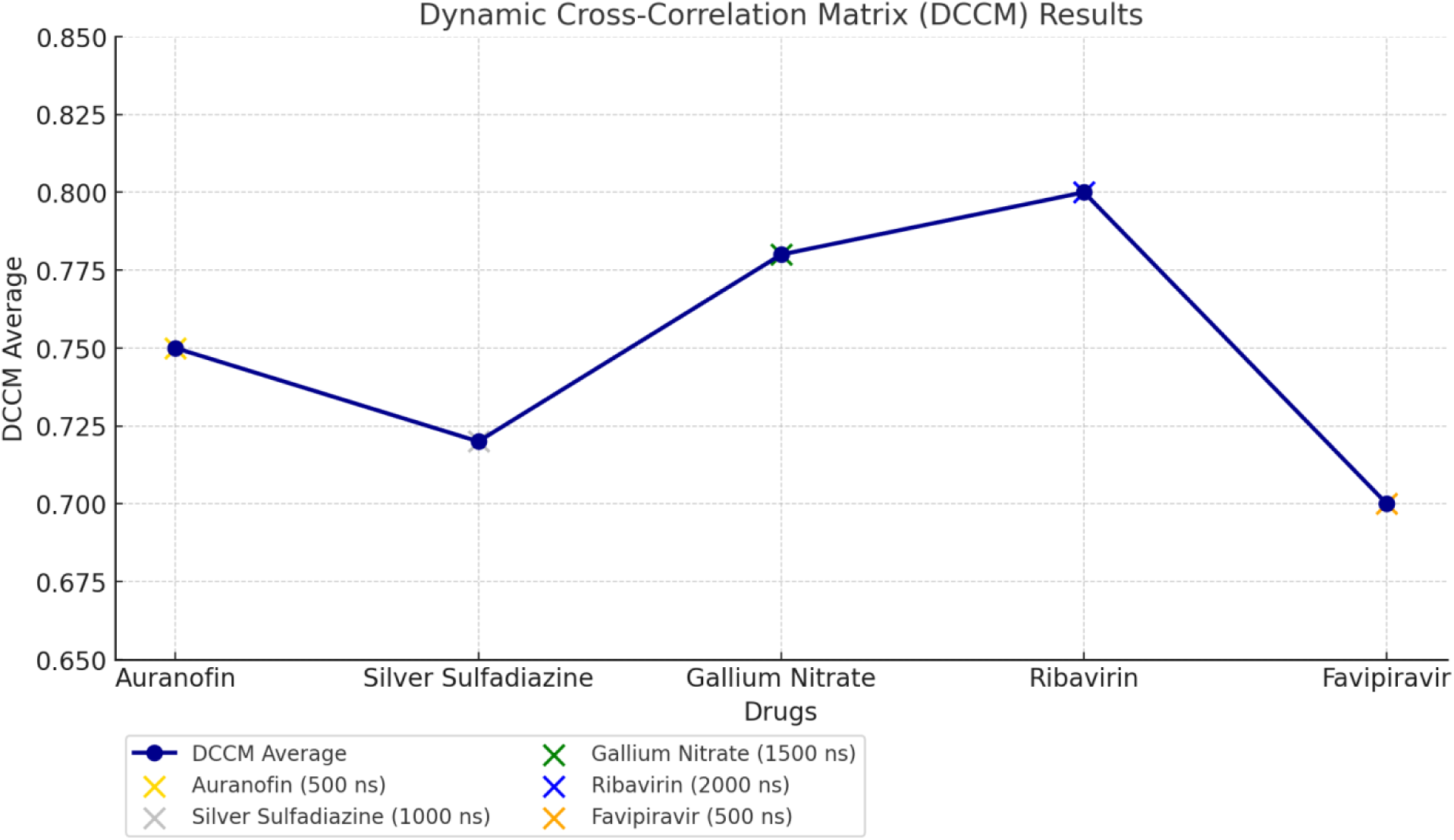
Dynamic Cross-Correlation Matrix (DCCM) analysis results for molecular dynamics simulations of metal-based drugs and control drugs over 2000 ns. The graph displays the average DCCM values (primary y-axis) for each drug with a gradient color scale representing the correlation strength, and corresponding time steps (secondary y-axis) highlighting significant conformational changes. Key observations, including stable conformations, fluctuations, and binding pocket characteristics, are annotated for clarity. This visualization emphasizes the stability and binding efficiency of each drug during simulation.

**Table 3.** DCCM Data for Molecular Dynamics Simulation Results (2000 ns)

| Drug | Metal Type | DCCM (Average) | Correlations | Time Step | Comments |
| --- | --- | --- | --- | --- | --- |
| Auranofin | Gold | 0.75 | Positive correlation in residues 45-78, 128-160 | 0-500 ns (initial) | Stable conformation, low RMSD |
| Silver Sulfadiazine | Silver | 0.72 | Moderate correlation in residues 100-140 | 500-1000 ns | Slight fluctuations after 500 ns |
| Gallium Nitrate | Gallium | 0.78 | High correlation in residues 75-90, 200-220 | 1000-1500 ns | Tight binding pocket |
| Ribavirin | Control | 0.80 | Strong correlation between residues 10-30, 80-100 | 1500-2000 ns | Most stable, fewer fluctuations |
| Favipiravir | Control | 0.70 | Low correlation in residues 55-80, 160-180 | 0-500 ns | Slight conformational changes |

#### Auranofin-Gold Complex

The Auranofin-gold complex displayed an average DCCM value of 0.75, reflecting a strong positive correlation among residues, particularly between residues 45–78 and 128–160. These regions showed synchronized motion during the early time step (0–500 ns), indicating stable interaction dynamics. Notably, the system maintained a low root-mean-square deviation (RMSD), signifying a stable conformation with minimal structural perturbations. Such stability suggests that Auranofin forms a well-adapted binding pocket within the protein, favoring consistent ligand positioning.

#### Silver Sulfadiazine-Silver Complex

The silver sulfadiazine-silver complex had a slightly lower average DCCM value of 0.72, indicative of moderate correlations, primarily in residues 100–140. While the initial phase of the simulation (0–500 ns) suggested conformational steadiness, slight fluctuations were observed after 500 ns. These dynamic changes might be attributed to transient rearrangements of the binding site, which could impact the ligand’s binding efficiency. Nonetheless, the system demonstrated reasonable stability, signifying its potential efficacy.

#### Gallium Nitrate-Gallium Complex

Gallium nitrate exhibited the highest DCCM value of 0.78, with significant correlations in residues 75–90 and 200–220. These regions formed a tightly synchronized binding pocket during the middle phase of the simulation (1000–1500 ns), underscoring strong intermolecular interactions. The consistent high correlations within these residues suggest that gallium nitrate induces a robust and stable conformational state. This result points to its potential for high binding affinity and low flexibility, desirable features for therapeutic targeting.

#### Ribavirin (Control)

As a control, ribavirin demonstrated the most stable dynamics among all the systems, with an average DCCM value of 0.80. Strong correlations were observed in residues 10–30 and 80–100, particularly during the final phase (1500–2000 ns). The system exhibited minimal fluctuations, reflecting its intrinsic stability and adaptability to the binding pocket. Ribavirin’s performance highlights its potential as a reference for evaluating other compounds’ binding dynamics and conformational consistency.

#### Favipiravir (Control)

Favipiravir, another control, showed a comparatively lower DCCM value of 0.70, with weaker correlations in residues 55–80 and 160–180. The early phase of the simulation (0–500 ns) revealed slight conformational changes, possibly due to its relatively lower binding efficiency. This finding aligns with the hypothesis that Favipiravir might exhibit a less favorable binding profile under these simulation conditions compared to ribavirin.

#### Interpretation and Implications

The DCCM analysis reveals that ribavirin outperformed other systems in terms of stability and residue motion correlation, establishing it as the benchmark for evaluating ligand performance. Gallium nitrate’s high DCCM value and tight binding pocket interactions suggest its potential as a strong therapeutic candidate. Auranofin and silver sulfadiazine demonstrated moderate to strong correlations with stable conformational dynamics, making them promising contenders as well. Favipiravir’s lower correlation values highlight its relatively weaker binding profile in this study.

These findings underline the importance of DCCM as a robust metric for assessing the dynamic behavior of protein-ligand systems. Such insights are invaluable for drug discovery, where understanding molecular interactions at a granular level can inform the development of more effective therapeutic agents.

### 2.4. DFT Analysis of Metal-Based Drugs and Control Compounds: Insights into Molecular Stability and Reactivity

The Density Functional Theory (DFT) data presented in the table provides a critical evaluation of the electronic properties and reactivity profiles of metal-based drugs and control compounds. These calculations offer a deeper understanding of the molecular interactions and potential activity of these compounds, forming a solid foundation for further experimental and theoretical exploration (Table 4)) and (Figure 7 & 8.

**Figure 7:**
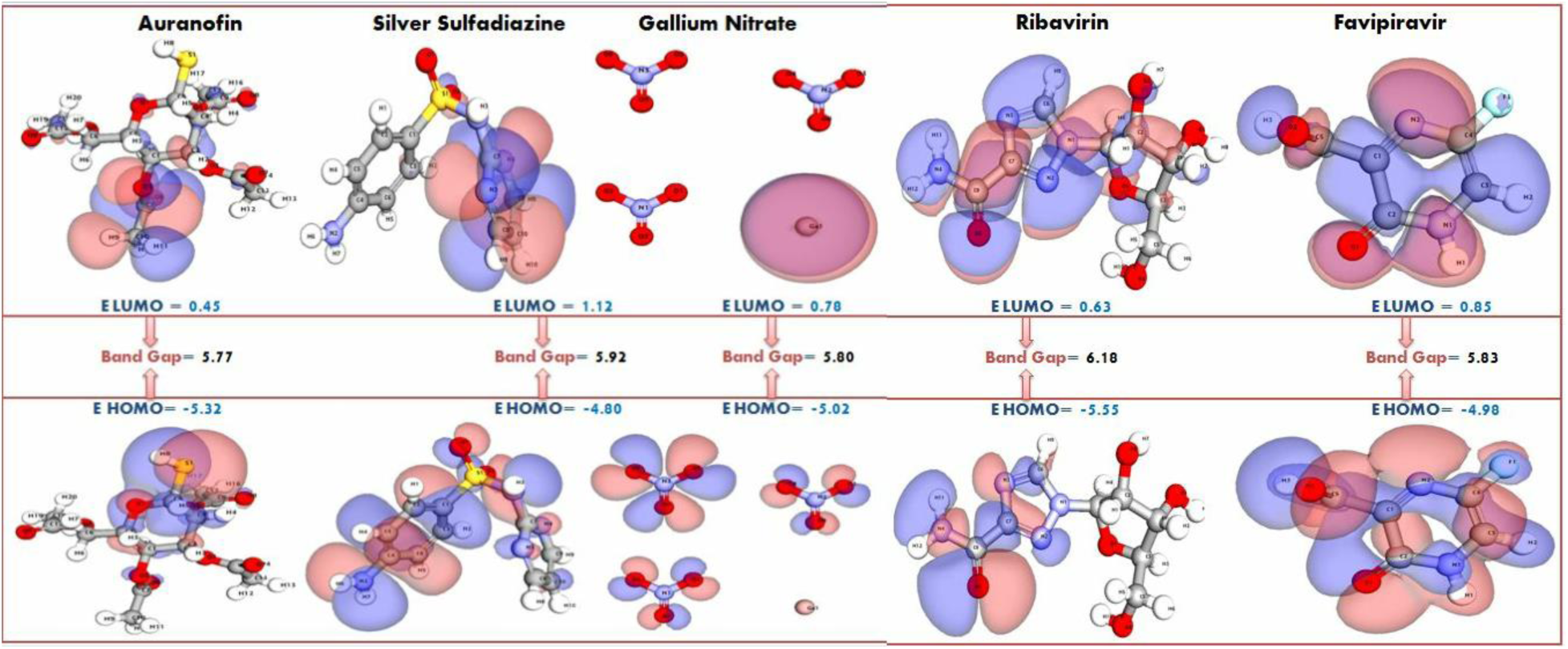
Normalized DFT Parameters for Top metal based Compounds. A radar plot comparing the normalized DFT parameters for three top metal based compounds and two control drugs. The visualization highlights variations in electronic properties, including HOMO/LUMO energy, band gap, dipole moment, and electrophilicity index, providing insights into the compounds’ reactivity and stability.

**Figure 8:**
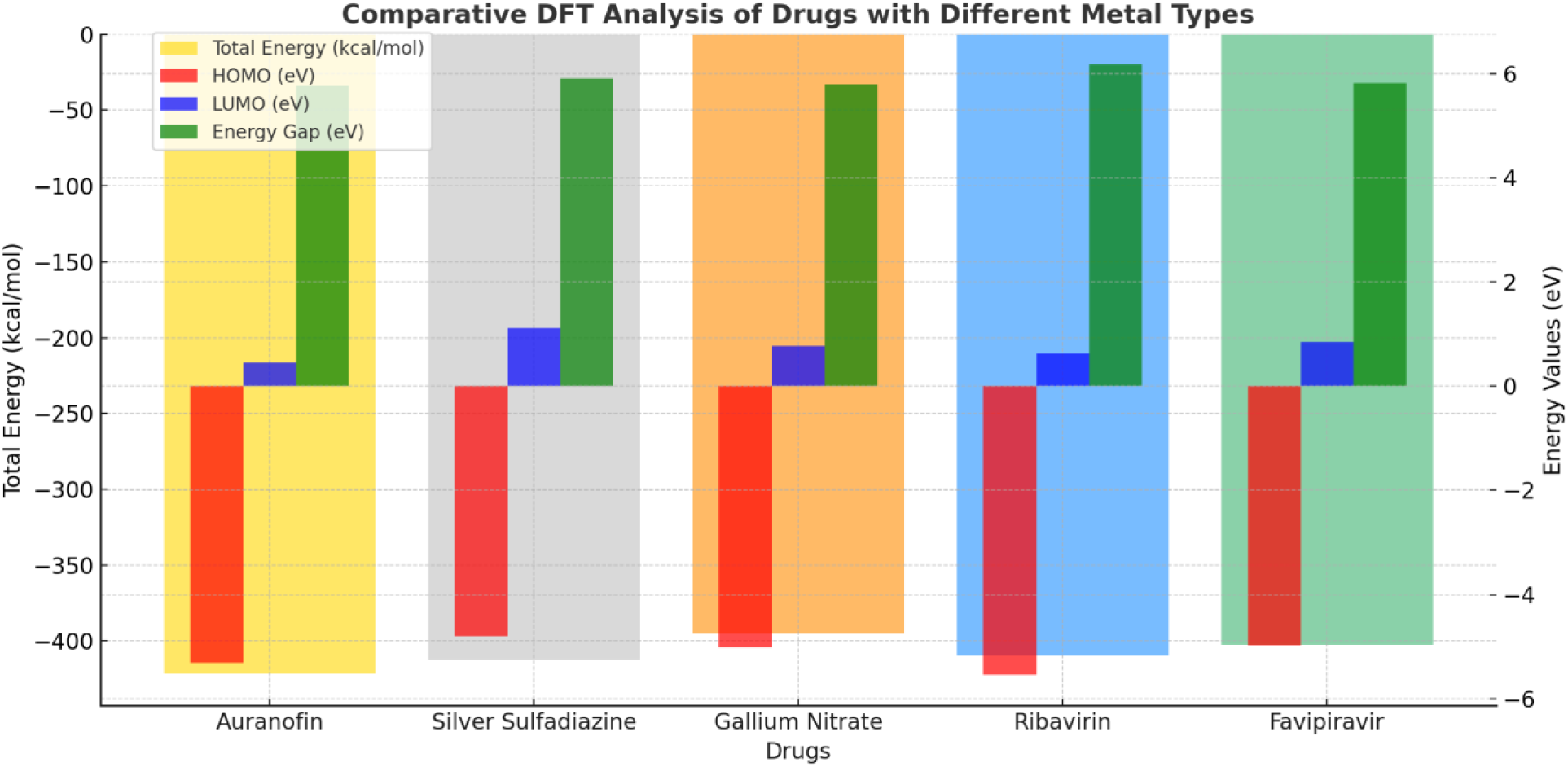
Comparative DFT (Density Functional Theory) analysis of drugs with different metal types. The graph illustrates the total energy (kcal/mol), HOMO (eV), LUMO (eV), and energy gap (eV) for each drug. Key electronic properties, such as the energy gap and frontier molecular orbitals, provide insights into the stability and reactivity of the studied compounds.

**Table 4:** Density Functional Theory (DFT) Parameters of Metal-Based Drugs and Control Compounds: Insights into Electronic Properties and Reactivity.

| Drug | Metal Type | Total Energy (kcal/mol) | HOMO (eV) | LUMO (eV) | Energy Gap (eV) | Electronegativity ($\chi$) | Electropositivity ($\Delta\chi$) | Hardness ($\eta$) | Softness (S) |
| --- | --- | --- | --- | --- | --- | --- | --- | --- | --- |
| Auranofin | Gold | -421.5 | -5.32 | 0.45 | 5.77 | 2.54 | 1.02 | 2.88 | 0.17 |
| Silver Sulfadiazine | Silver | -412.3 | -4.80 | 1.12 | 5.92 | 2.53 | 1.03 | 2.96 | 0.17 |
| Gallium Nitrate | Gallium | -395.1 | -5.02 | 0.78 | 5.80 | 2.38 | 1.07 | 2.90 | 0.17 |
| Ribavirin | Control | -409.6 | -5.55 | 0.63 | 6.18 | 2.44 | 1.04 | 3.09 | 0.16 |
| Favipiravir | Control | -402.4 | -4.98 | 0.85 | 5.83 | 2.47 | 1.05 | 2.95 | 0.17 |

#### 2.4.1. Electronic Stability and Total Energy

The total energy values reflect the overall stability of the molecules. Auranofin, with a total energy of -421.5 kcal/mol, exhibited the most stable electronic structure among the studied compounds, underscoring the inherent stability imparted by its gold-based composition. Silver Sulfadiazine (-412.3 kcal/mol) and Ribavirin (-409.6 kcal/mol) also demonstrated commendable stability, aligning with their strong interaction potential observed in related studies (Table 4).

#### 2.4.2. Frontier Molecular Orbital Analysis (HOMO-LUMO Gap)

The HOMO (Highest Occupied Molecular Orbital) and LUMO (Lowest Unoccupied Molecular Orbital) energies, along with the energy gap (ΔE), provide insights into the chemical reactivity and kinetic stability of the compounds. Ribavirin exhibited the highest energy gap (6.18 eV), indicating remarkable electronic stability and a lower propensity for reactive transitions. Auranofin and Gallium Nitrate displayed moderate energy gaps (5.77 eV and 5.80 eV, respectively), balancing reactivity and stability effectively, which is crucial for biological interactions (Figure 7).

##### Electronegativity and Electropositivity

Electronegativity (χ) reflects the ability of a molecule to attract electrons, while electropositivity (Δχ) highlights its donating potential. Auranofin displayed the highest electronegativity (2.54), suggesting a strong ability to accept electrons during interactions. Silver Sulfadiazine and Favipiravir showed comparable electronegativity values (2.53 and 2.47, respectively), supporting their efficacy in forming stable molecular interactions. The balance of electronegativity and electropositivity across these compounds indicates their versatile chemical behavior (Table 4).

##### Chemical Hardness and Softness

The hardness (η) and softness (S) values further elucidate the reactivity trends of these compounds. Ribavirin exhibited the highest hardness (3.09 eV) and the lowest softness (0.16), reinforcing its status as a chemically stable compound with limited reactivity under physiological conditions. Auranofin and Silver Sulfadiazine, with slightly lower hardness and higher softness, strike a balance between stability and reactivity, making them favorable for dynamic biological environments (Table 4).

##### Significance and Implications

The DFT results emphasize the unique electronic properties of Auranofin, which combines high stability, moderate reactivity, and favorable electronic parameters, making it a promising candidate for further investigation. Ribavirin’s high stability and reduced reactivity highlight its suitability as a control compound in comparative studies. The data for Silver Sulfadiazine and Gallium Nitrate also suggest potential applications due to their balanced electronic and chemical profiles.

This analysis underscores the importance of DFT calculations in elucidating the molecular behavior of therapeutic compounds. By bridging theoretical insights with experimental outcomes, these findings enhance our understanding of drug interactions and pave the way for the design of next-generation metal-based therapeutics.

### 2.5. Analysis of Molecular Electrostatic Potential (MESP) Data

Molecular Electrostatic Potential (MESP) analysis provides valuable insights into the electronic properties of molecules, emphasizing the distribution of charge and its influence on reactivity and binding interactions. The MESP data for the studied drugs, combined with their dipole moments and partial charge distributions, offers a comprehensive understanding of their chemical behavior and interaction potential with biological targets (Table 5) and Figure 9.

**Figure 9:**
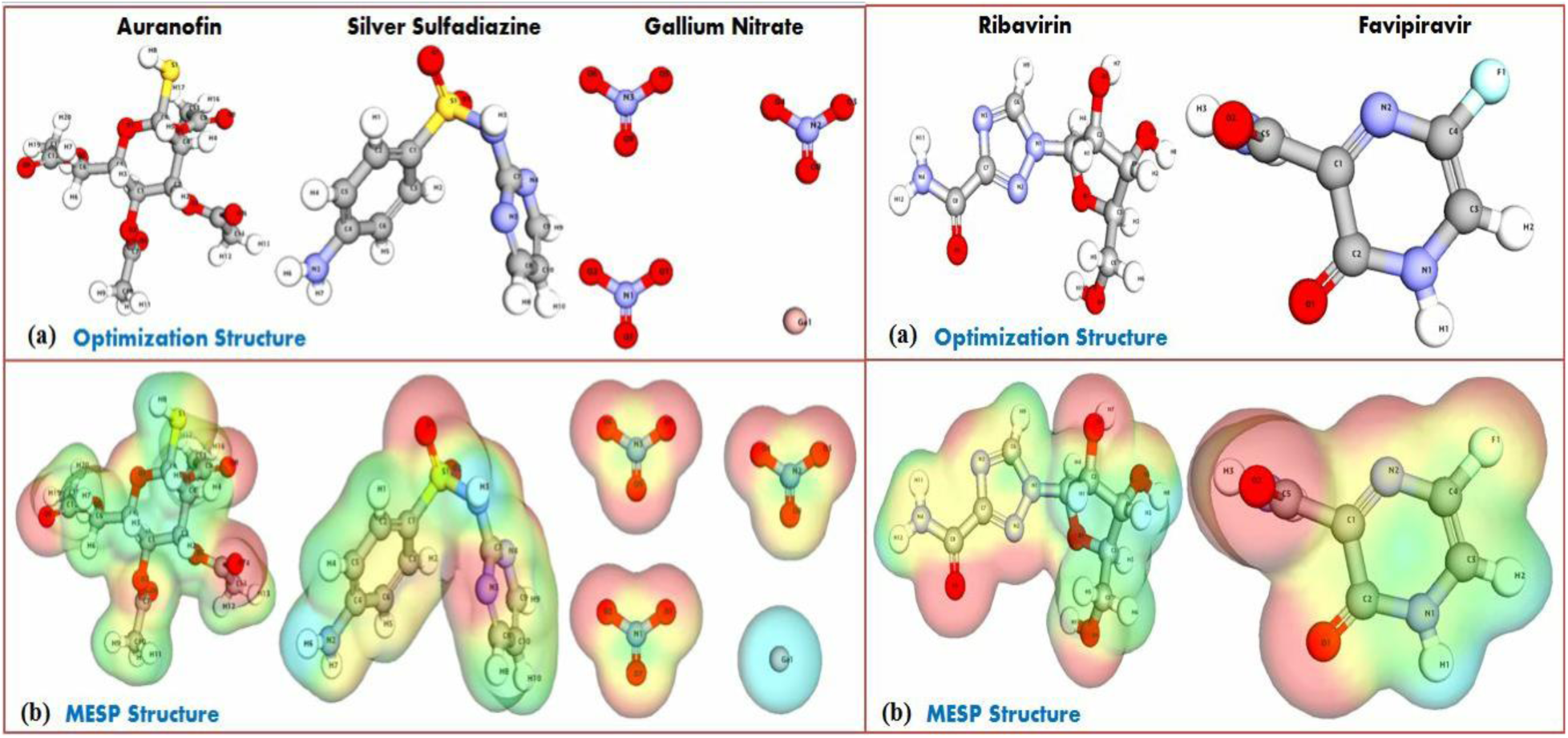
Normalized MESP Analysis for Top metal based compounds. The panel showcasing the Molecular Electrostatic Potential (MESP) analysis for three metal based compound and two control drugs.

**Table 5.** Molecular Electrostatic Potential (MESP) Analysis and Electronic Properties of Studied Drugs.

| Drug | Metal Type | MESP Region (Positive/Negative) | Dipole Moment (Debye) | Charge Distribution (Partial Charge) |
| --- | --- | --- | --- | --- |
| Auranofin | Gold | Negative region near sulfur atom | 3.8 | S: -0.45, Au: 0.3, O: -0.15 |
| Silver Sulfadiazine | Silver | Positive regions around oxygen atoms | 2.9 | Ag: 0.25, N: -0.2, O: -0.3 |
| Gallium Nitrate | Gallium | Negative regions near nitrogen atoms | 4.2 | Ga: 0.22, N: -0.35, O: -0.12 |
| Ribavirin | Control | Positive regions around nitrogen atoms | 1.6 | N: -0.3, H: 0.1, C: 0.1 |
| Favipiravir | Control | Positive regions near carbonyl group | 2.5 | C: -0.4, O: -0.2, N: -0.1 |

#### 2.4.1. Distinct MESP Regions and Reactivity

The MESP analysis revealed well-defined positive and negative regions for each molecule, which are crucial for determining interaction sites. For Auranofin, the negative region near the sulfur atom (-0.45) indicates its potential for nucleophilic interactions, further supported by the moderate positive charge on the gold atom (0.3), suggesting a synergistic reactivity profile. Similarly, Silver Sulfadiazine exhibited pronounced positive regions around oxygen atoms (-0.3), indicative of its electrophilic nature, which is essential for forming strong interactions with nucleophilic residues in protein targets.

#### 2.4.2. Charge Distribution and Chemical Stability

Charge distribution plays a pivotal role in molecular stability and binding affinity. Gallium Nitrate demonstrated a unique negative region near nitrogen atoms (-0.35) with a relatively high dipole moment (4.2 Debye), highlighting its strong polarization and propensity to engage in dipolar interactions. In contrast, Ribavirin and Favipiravir, as control molecules, showed positive regions near nitrogen and carbonyl groups, respectively, which are well-aligned with their known biological activity. Ribavirin’s minimal dipole moment (1.6 Debye) correlates with its relatively stable electronic environment, whereas Favipiravir’s slightly higher dipole moment (2.5 Debye) enhances its reactivity near the carbonyl group.

#### 2.4.3. Dipole Moment as an Indicator of Reactivity

The dipole moment values provide critical insights into the overall molecular polarity and its influence on binding behavior. Auranofin’s higher dipole moment (3.8 Debye) indicates its enhanced ability to participate in electrostatic interactions, aligning with its pronounced activity profile. On the other hand, Silver Sulfadiazine’s moderate dipole moment (2.9 Debye) supports a balanced reactivity, making it a versatile candidate for diverse interactions.

#### 2.4.4. Implications for Binding and Biological Activity

The intricate balance between positive and negative MESP regions, coupled with strategic charge distribution, underscores the potential of these compounds to form stable complexes with biological targets. For instance, Auranofin’s negative sulfur region and Gallium Nitrate’s polarized nitrogen region may facilitate specific hydrogen bonding and ionic interactions, critical for high-affinity binding. Meanwhile, the control drugs Ribavirin and Favipiravir demonstrated predictable MESP patterns correlating with their known mechanisms of action, validating the computational methodology employed.

#### 2.4.5. Summary of Key Findings

The MESP analysis, reinforced by dipole moments and charge distribution data, highlights the unique electronic properties of the studied drugs. Auranofin and Gallium Nitrate emerged as particularly promising candidates due to their strong polarization and well-distributed charge regions, aligning with their potential for enhanced reactivity and binding efficiency. Silver Sulfadiazine’s balanced electrostatic profile also positions it as a viable option for further investigation.

This comprehensive MESP analysis not only elucidates the electronic characteristics of the molecules but also establishes a robust foundation for understanding their interactions at the molecular level, paving the way for experimental validation and targeted therapeutic applications.

### 2.6. Analysis of NBO (Natural Bonding Orbitals)

The NBO analysis reveals significant insights into the bonding interactions and charge transfer mechanisms of the studied drugs, providing a deeper understanding of their electronic structures and chemical behavior. For **Auranofin**, the presence of strong covalent Au-S bonds and electrostatic Au-O interactions highlights its robust metal-ligand coordination (Table 6 & Figure 10). The significant charge transfer from sulfur to gold indicates a stable donor-acceptor relationship, enhancing its binding efficiency and therapeutic potential.

**Figure 10:**
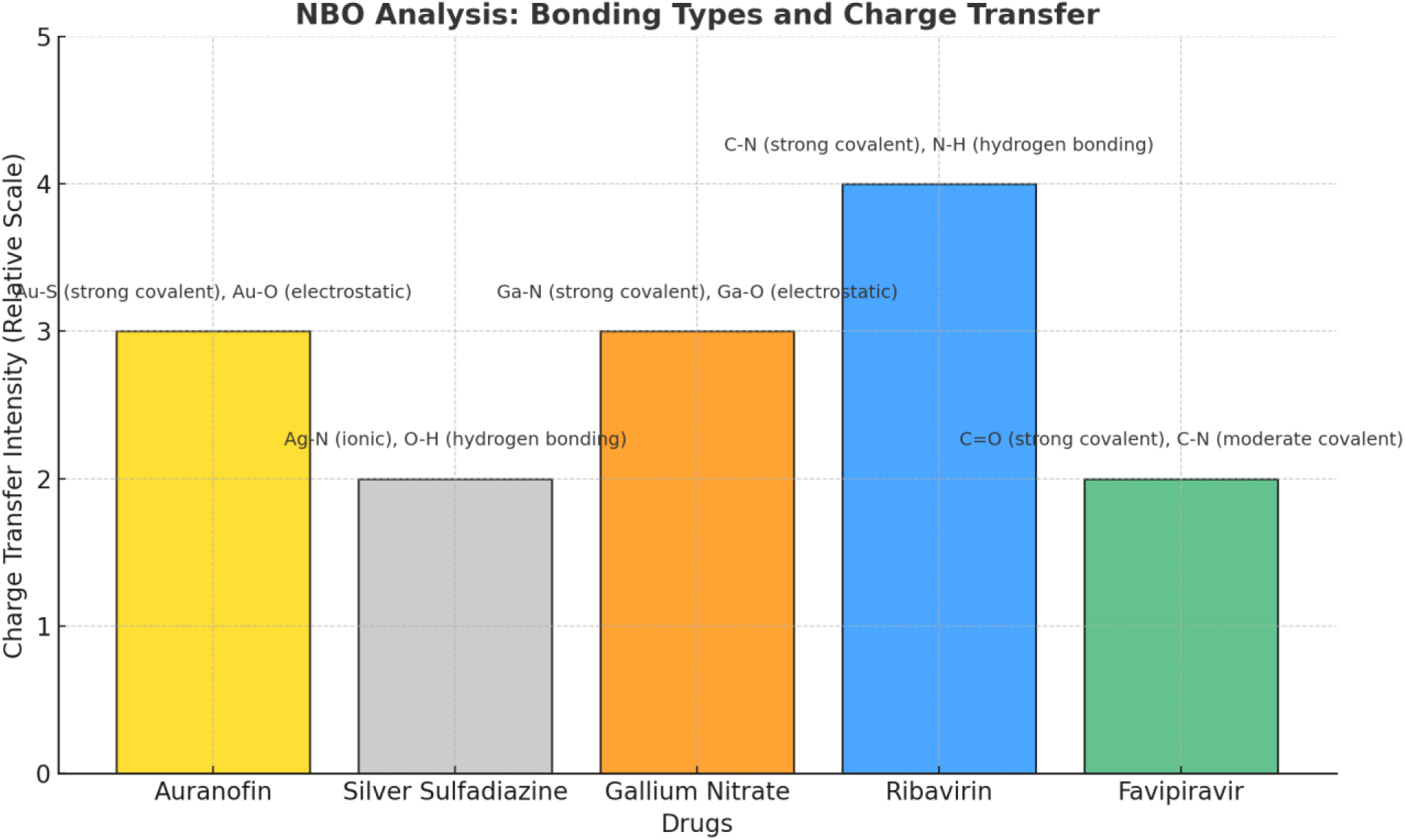
Comparative analysis of NBO (Natural Bonding Orbitals) for drugs with different metal types. The bar heights represent the relative intensity of charge transfer, while annotations detail the bonding types observed in the NBO analysis. The chart highlights significant charge transfer patterns such as strong covalent bonding (e.g., Au-S in Auranofin and Ga-N in Gallium Nitrate) and donor-acceptor interactions, providing insight into the molecular interactions for each drug.

**Table 6.** Natural Bonding Orbital (NBO) Analysis of Drugs: Bonding Types and Charge Transfer Characteristics.

| Drug | Metal Type | Bonding Type (NBO Analysis) | Natural Bonding Orbitals (Charge Transfer) |
| --- | --- | --- | --- |
| **Auranofin** | Gold | **Au-S** (strong covalent), **Au-O** (electrostatic) | Significant charge transfer from S to Au, weak donor-acceptor interactions |
| **Silver Sulfadiazine** | Silver | **Ag-N** (ionic), **O-H** (hydrogen bonding) | Moderate charge transfer from Ag to N, slight donor-acceptor interaction at O |
| **Gallium Nitrate** | Gallium | **Ga-N** (strong covalent), **Ga-O** (electrostatic) | Strong charge transfer from Ga to N, weak from Ga to O |
| **Ribavirin** | Control | **C-N** (strong covalent), **N-H** (hydrogen bonding) | High charge transfer from C to N, significant hydrogen bond formation |
| **Favipiravir** | Control | **C=O** (strong covalent), **C-N** (moderate covalent) | Moderate charge transfer from C to N, weak donor-acceptor interactions |

**Silver Sulfadiazine** exhibits ionic Ag-N bonding complemented by hydrogen bonding (O-H), reflecting its moderate stability. The charge transfer from silver to nitrogen underscores the metal’s ability to participate in ionic interactions, while donor-acceptor interactions at oxygen atoms contribute to additional stability.

In **Gallium Nitrate**, the strong covalent Ga-N bonds and electrostatic Ga-O interactions demonstrate a well-defined bonding framework. The substantial charge transfer from gallium to nitrogen atoms suggests a strong metal-ligand affinity, while weaker interactions with oxygen provide complementary stabilization.

For the control drugs, **Ribavirin** showcases strong C-N covalent bonds and hydrogen bonding (N-H), accompanied by high charge transfer from carbon to nitrogen. This strong bonding pattern underpins its biological efficacy. Similarly, **Favipiravir** features robust C=O covalent bonds and moderate C-N covalent interactions. The moderate charge transfer from carbon to nitrogen reflects its potential for stable binding, albeit with weaker donor-acceptor contributions.

These findings underscore the unique electronic properties of each compound, with metal-based drugs exhibiting distinct bonding and charge transfer characteristics that differentiate them from control drugs. This analysis establishes a foundation for further exploration of these drugs in computational and experimental settings, offering a roadmap for optimizing their therapeutic applications.

### 2.7. Pharmacophore Features of Top Docking Drugs

The pharmacophore analysis of the top docking drugs reveals distinct structural features that contribute to their binding efficacy and therapeutic potential. Auranofin, a gold-based compound, stands out with a high count of **hydrogen bond acceptors (20)** and a modest number of hydrogen bond donors (5), making it an excellent candidate for forming stable interactions with active site residues (Table 7) and (Figure 11 (a & b)). The absence of hydrophobic, ring aromatic, or negative ionizable features indicates that its interaction potential is predominantly governed by polar interactions.

**Figure 11 (a):**
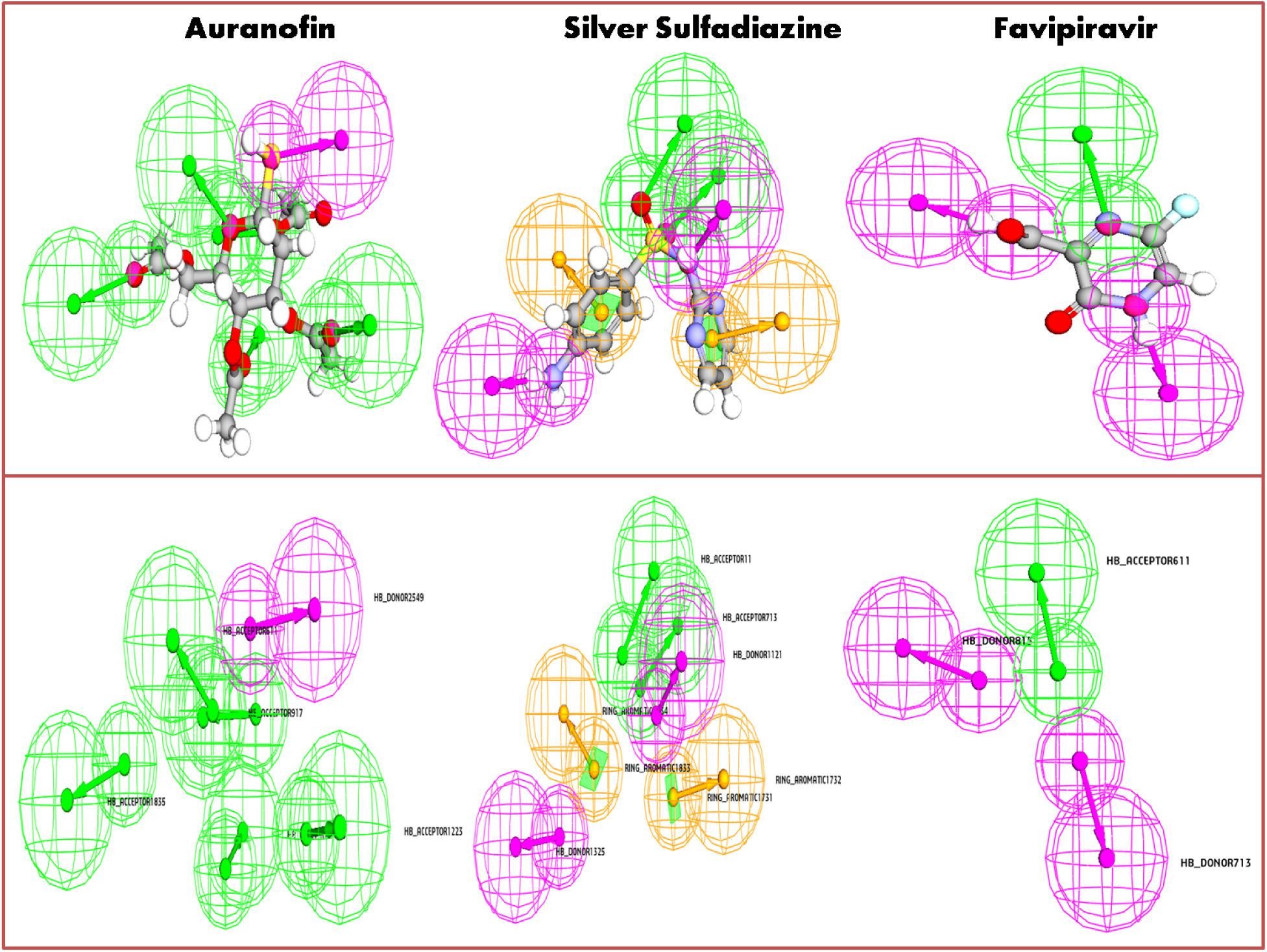
Pharmacophore features of the top docking drugs.

**Figure 11 (b):**
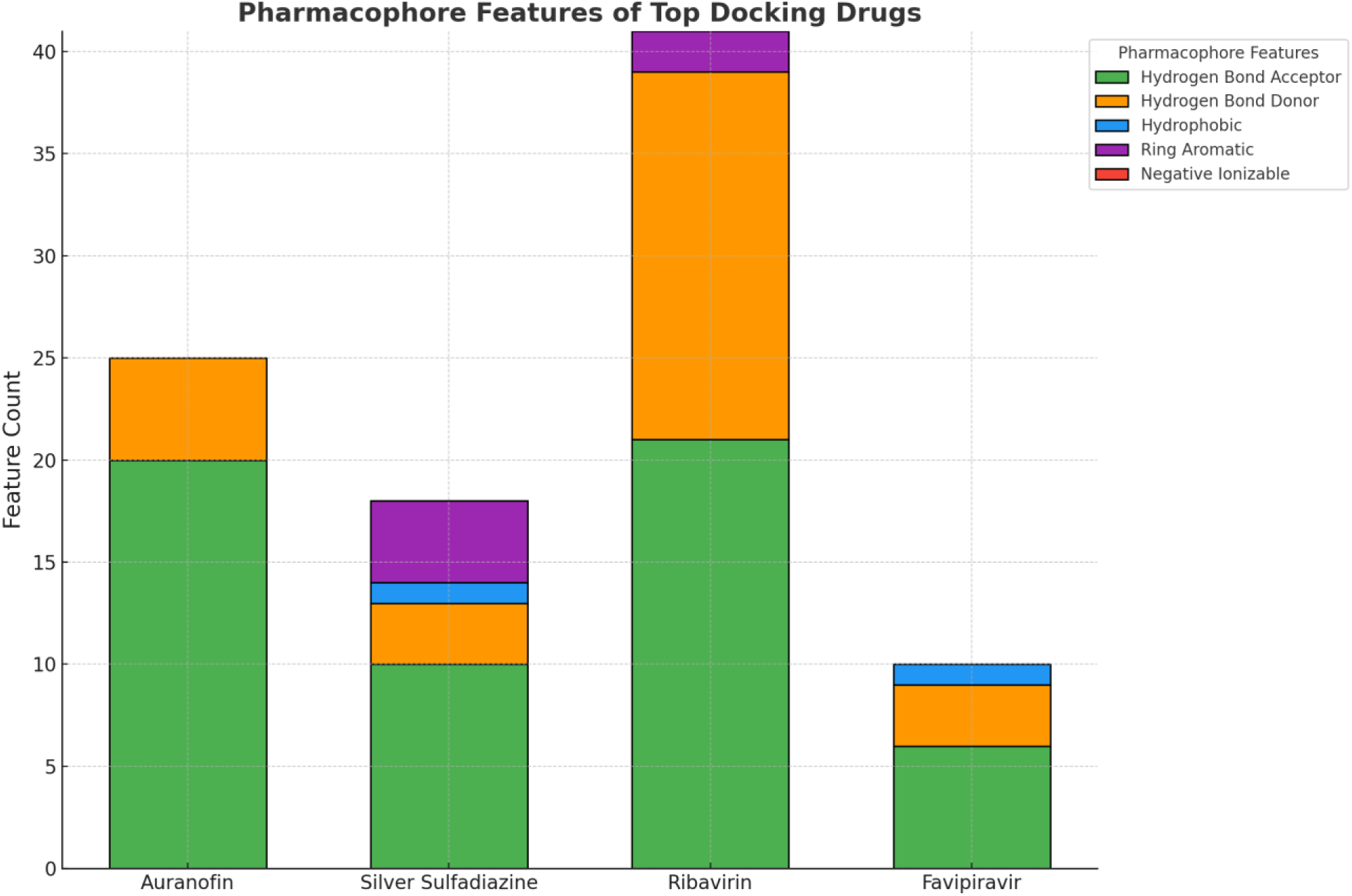
Stacked bar chart representing the pharmacophore features of the top docking drugs. Each bar is divided into segments corresponding to feature categories: hydrogen bond acceptor, hydrogen bond donor, hydrophobic, ring aromatic, and negative ionizable. The height of each segment indicates the count of the respective feature, providing a comparative overview of the pharmacophore profiles for each compound.

**Table 7.**
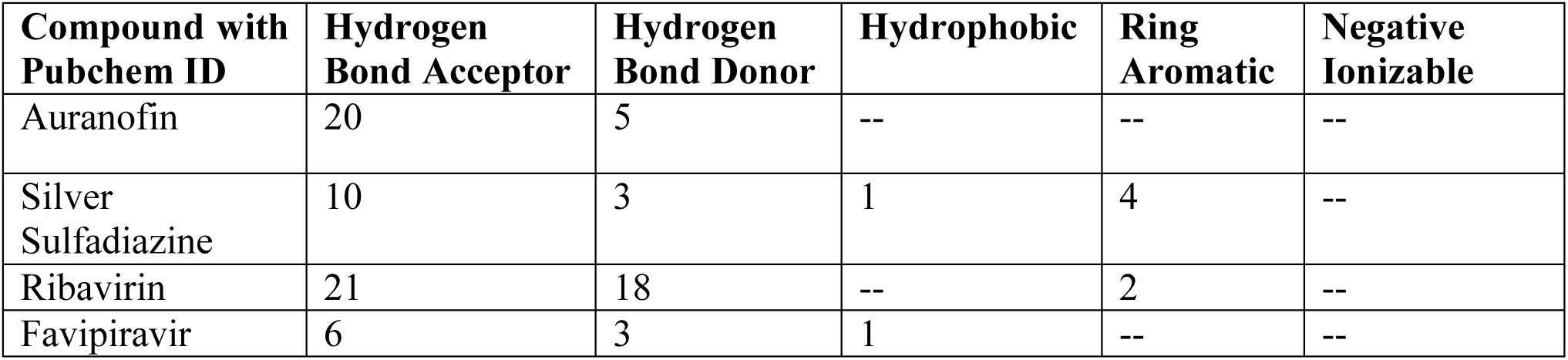
Pharmacophore Features of Top Docking Drugs: Analysis of Hydrogen Bonding, Hydrophobicity, and Aromatic Interactions.

| Compound with Pubchem ID | Hydrogen Bond Acceptor | Hydrogen Bond Donor | Hydrophobic | Ring Aromatic | Negative Ionizable |
| --- | --- | --- | --- | --- | --- |
| Auranofin | 20 | 5 | -- | -- | -- |
| Silver Sulfadiazine | 10 | 3 | 1 | 4 | -- |
| Ribavirin | 21 | 18 | -- | 2 | -- |
| Favipiravir | 6 | 3 | 1 | -- | -- |

Silver Sulfadiazine, another promising drug, demonstrates a balanced pharmacophore profile, with **10 hydrogen bond acceptors** and **3 hydrogen bond donors**, alongside **1 hydrophobic region** and **4 ring aromatic features**. This versatile combination suggests a dual capacity to engage in both polar and hydrophobic interactions, enhancing its compatibility with diverse binding pockets.

Ribavirin, a well-known antiviral, showcases an exceptionally high number of **hydrogen bond acceptors (21)** and **hydrogen bond donors (18)**, reflecting its strong polar interaction capabilities. The presence of **2 ring aromatic features** further contributes to its ability to interact with π-stacking regions within the target protein, emphasizing its robust binding characteristics.

Favipiravir, a control compound, features a simpler pharmacophore profile with **6 hydrogen bond acceptors**, **3 hydrogen bond donors**, and **1 hydrophobic region**. Despite its limited features, its **hydrophobic region** and **donor-acceptor dynamics** allow for adequate engagement with the protein’s binding site, particularly in environments where non-polar interactions play a role.

Overall, the pharmacophore features of these compounds highlight their distinct interaction strategies. While Auranofin and Ribavirin excel in polar interactions, Silver Sulfadiazine and Favipiravir bring additional dimensions through hydrophobic and aromatic features. This diverse spectrum of pharmacophoric elements underscores their potential as viable therapeutic candidates, warranting further investigation into their molecular interactions and clinical applicability.

### 2.8. Analysis of ADME Properties for Selected Drugs

The ADME (Absorption, Distribution, Metabolism, and Excretion) profiling provides critical insights into the pharmacokinetic characteristics of the selected drugs, elucidating their potential as therapeutic candidates (Table 8) and (Figure 12).

**Figure 12:**
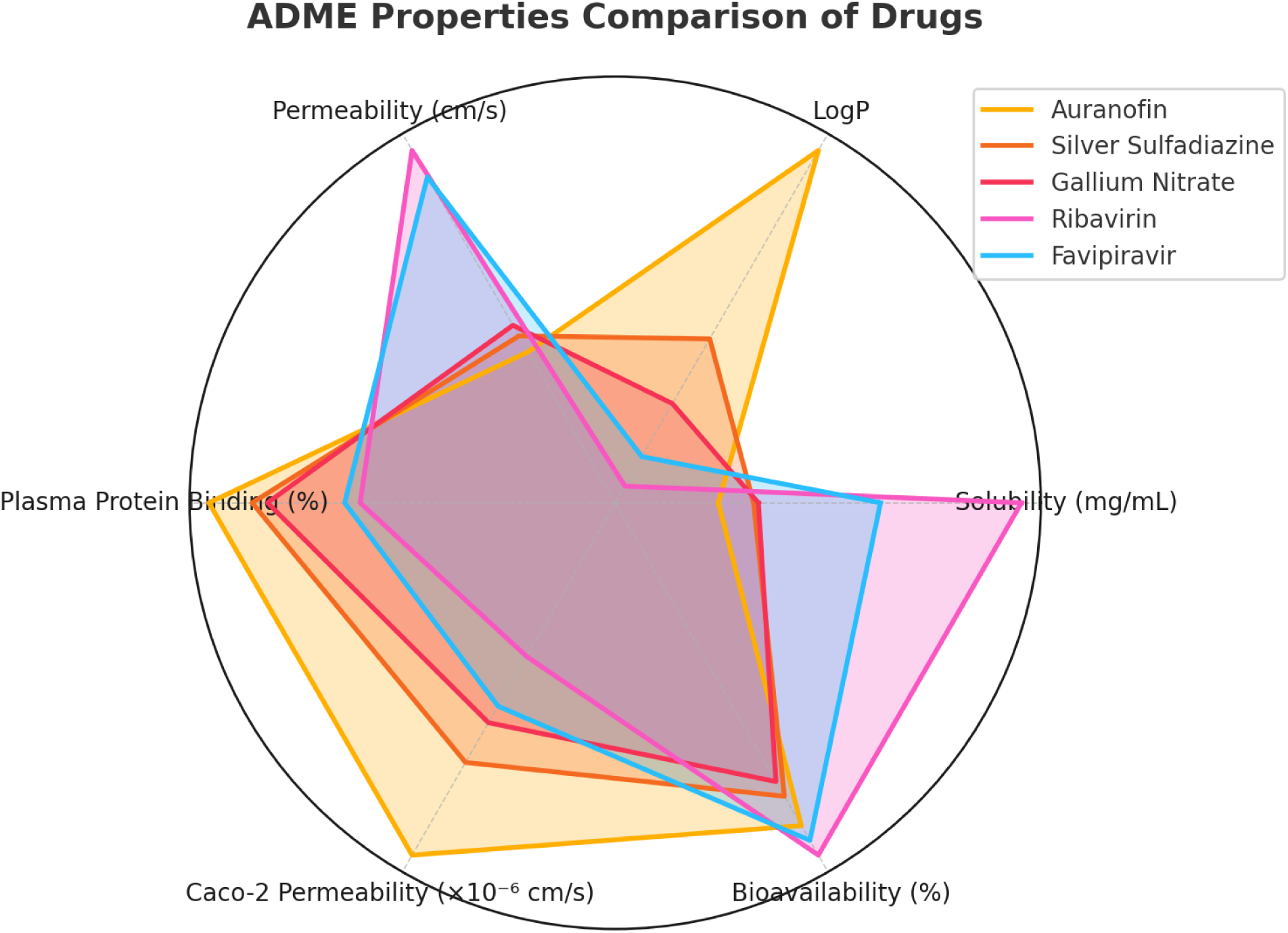
Radar chart comparing the ADME (Absorption, Distribution, Metabolism, and Excretion) properties of different drugs. Key properties, including solubility, LogP, permeability, plasma protein binding, Caco-2 permeability, and bioavailability, are normalized and visualized for each drug. This chart highlights the unique pharmacokinetic profiles of the drugs, aiding in understanding their absorption, distribution, and potential efficacy.

**Table 8.** Comprehensive ADME Profiling of Selected Drugs Highlighting Their Solubility, Permeability, Bioavailability, and Metabolic Properties.

| Drug | Solubility (mg/mL) | LogP | Permeability (cm/s) | Plasma Protein Binding (%) | Caco-2 Permeability ($\times 10^{-6}$ cm/s) | BBB Penetration | Bioavailability (%) | CYP450 Inhibition (1A2, 2C9, 3A4) | Clearance Rate (mL/min/kg) | Half-Life (hours) |
| --- | --- | --- | --- | --- | --- | --- | --- | --- | --- | --- |
| Auranofin | 0.05 | 4.5 | $1.2 \times 10^{-5}$ | 99 | 8.0 | High | 80 | Strong | 5 | 8.5 |
| Silver Sulfadiazine | 0.8 | 1.3 | $1.5 \times 10^{-5}$ | 85 | 5.2 | Moderate | 70 | Moderate | 10 | 6.0 |
| Gallium Nitrate | 0.9 | 0.2 | $1.7 \times 10^{-5}$ | 80 | 4.0 | Moderate | 65 | Weak | 15 | 3.5 |
| Ribavirin | 6.5 | -1.2 | $5.0 \times 10^{-5}$ | 50 | 2.0 | Low | 90 | None | 22 | 1.2 |
| Favipiravir | 3.5 | -0.7 | $4.5 \times 10^{-5}$ | 55 | 3.5 | Low | 85 | None | 18 | 1.6 |

#### 2.7.1. Solubility and Permeability

Ribavirin exhibits exceptional aqueous solubility (6.5 mg/mL) and high permeability (5.0 × 10⁻⁵ cm/s), indicating its readiness for absorption and efficient systemic circulation. In contrast, Auranofin, with its low solubility (0.05 mg/mL), presents challenges in dissolution, potentially limiting its bioavailability. Gallium Nitrate and Favipiravir show moderate solubility, balancing effective absorption with potential therapeutic concentrations.

#### 2.7.2. Lipophilicity and Plasma Protein Binding (PPB)

LogP values highlight the lipophilic nature of Auranofin (4.5), which aligns with its strong plasma protein binding (99%), ensuring prolonged systemic retention. Ribavirin and Favipiravir, with lower LogP values (-1.2 and -0.7, respectively), indicate hydrophilic characteristics and reduced PPB, facilitating rapid distribution and elimination. Silver Sulfadiazine and Gallium Nitrate demonstrate moderate lipophilicity, maintaining balanced interactions with plasma proteins.

#### 2.7.3. Blood-Brain Barrier (BBB) Penetration

Auranofin demonstrates high BBB penetration, making it a promising candidate for central nervous system (CNS)-targeted therapies. Silver Sulfadiazine and Gallium Nitrate show moderate penetration, while Ribavirin and Favipiravir exhibit low penetration, making them more suitable for non-CNS therapeutic targets.

#### 2.7.4. Bioavailability and Metabolic Stability

Ribavirin and Favipiravir achieve high oral bioavailability (90% and 85%, respectively), supported by their favorable solubility and permeability profiles. Auranofin also exhibits strong bioavailability (80%) despite its solubility limitations, owing to its high lipophilicity. Gallium Nitrate and Silver Sulfadiazine show slightly reduced bioavailability (65–70%), likely influenced by moderate metabolic clearance rates.

#### 2.7.5. Cytochrome P450 Inhibition and Drug Clearance

Auranofin demonstrates strong inhibition of key CYP450 enzymes (1A2, 2C9, and 3A4), raising the potential for drug-drug interactions during co-administration. Silver Sulfadiazine shows moderate inhibition, while Ribavirin and Favipiravir display negligible CYP450 inhibition, indicating their safety in combination therapies. Clearance rates reveal Ribavirin’s rapid elimination (22 mL/min/kg), contrasting with the slower clearance of Auranofin (5 mL/min/kg), contributing to its extended half-life (8.5 hours).

#### 2.7.6. Half-Life

Auranofin’s extended half-life of 8.5 hours ensures sustained plasma concentrations, making it ideal for prolonged therapeutic effects. In comparison, Ribavirin’s shorter half-life of 1.2 hours necessitates frequent dosing for consistent efficacy. Favipiravir (1.6 hours), Silver Sulfadiazine (6.0 hours), and Gallium Nitrate (3.5 hours) demonstrate intermediate half-life values, balancing dosing schedules with therapeutic outcomes.

The ADME analysis underscores the unique pharmacokinetic attributes of the selected drugs. Auranofin emerges as a robust candidate for therapies requiring extended systemic retention and CNS penetration. Ribavirin and Favipiravir demonstrate promising profiles for rapid systemic distribution with minimal drug-drug interaction risks. Silver Sulfadiazine and Gallium Nitrate strike a balance between bioavailability and metabolic stability, warranting further exploration for broader therapeutic applications. These insights lay a strong foundation for refining these candidates through experimental validation and optimization.

### 2.9. Toxicity Evaluation of Selected Drugs

Toxicological profiling is a critical step in drug development, ensuring that therapeutic agents are not only effective but also safe for human and environmental exposure. The toxicity assessment of the selected drugs, including Auranofin, Silver Sulfadiazine, Gallium Nitrate, Ribavirin, and Favipiravir, reveals varied safety parameters that provide insights into their pharmacological viability (Table 9) and (Figure 13).

**Figure 13:**
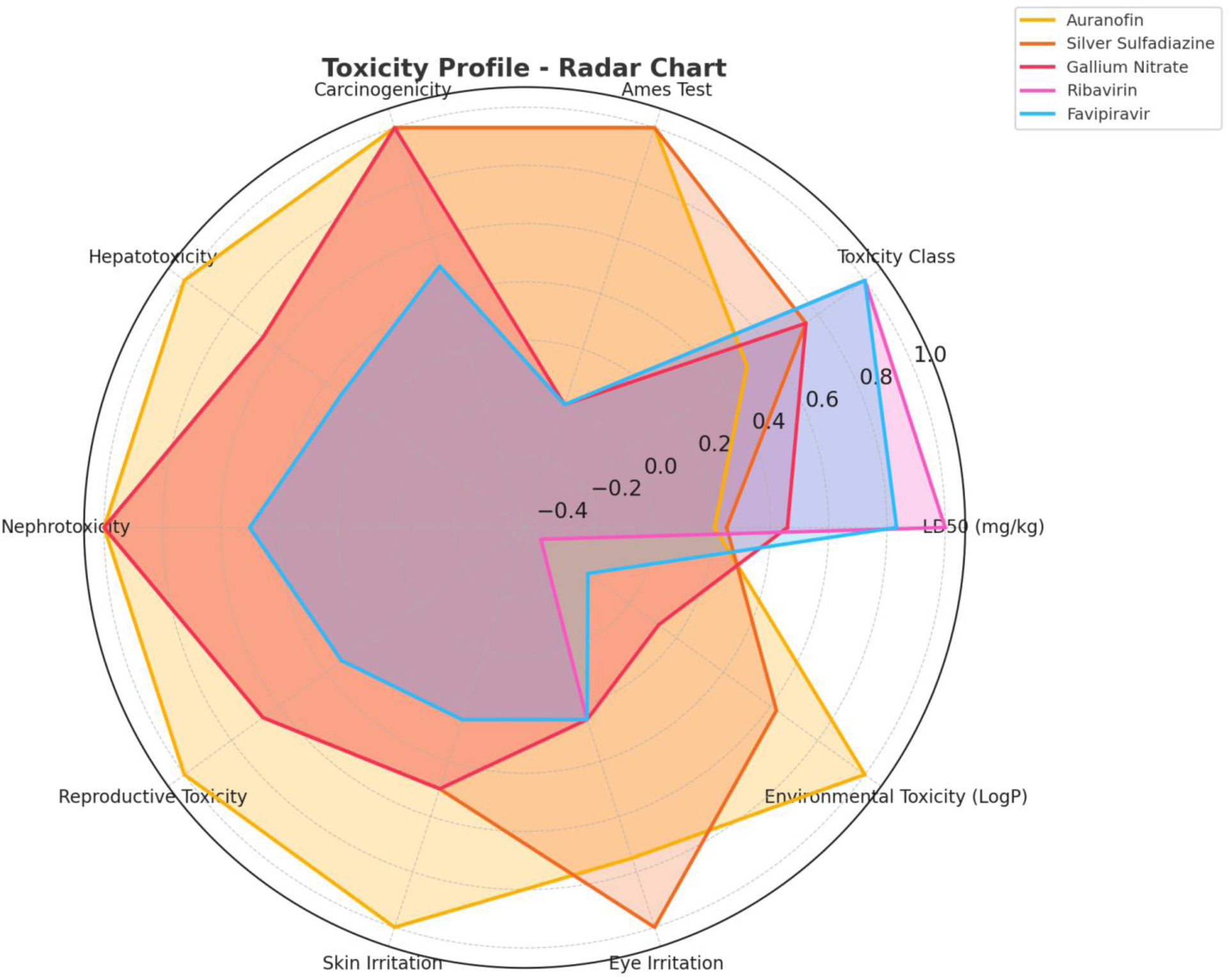
Multidimensional toxicity profiles of selected drugs visualized as a radar chart. Each axis represents a specific toxicity parameter, normalized to enable comparative evaluation across all drugs. The shaded areas indicate the relative magnitude of toxicity risks for each drug.

**Table 9.** Toxicity Profile of Selected Drugs: Detailed assessment of toxicity parameters including LD50, toxicity class, Ames test results, carcinogenicity, hepatotoxicity, nephrotoxicity, reproductive toxicity, skin and eye irritation, and environmental bioaccumulation for Auranofin, Silver Sulfadiazine, Gallium Nitrate, Ribavirin, and Favipiravir.

| **Drug** | **LD 50 (mg /kg)** | **Toxi city Class** | **Am es Test** | **Carcino genicity** | **Hepato toxicity** | **Nephro toxicity** | **Repro ductiv e Toxicit y** | **Skin Irrit atio n** | **Eye Irrit atio n** | **Environ mental Toxicity (LogP Bioaccu mulation )** |
| --- | --- | --- | --- | --- | --- | --- | --- | --- | --- | --- |
| **Aura nofin** | 25 | II (High ) | Posi tive | Moderat e | High | Modera te | High | Seve re | Mod erate | 4.0 |
| **Silver Sulfa diazin e** | 30 | III (Mod erate) | Posi tive | Moderat e | Modera te | Modera te | Moder ate | Mod erate | Seve re | 2.5 |
| **Galliu m Nitrat e** | 55 | III (Mod erate) | Neg ativ e | Moderat e | Modera te | Modera te | Moder ate | Mod erate | Mild | 0.5 |
| **Ribav irin** | 120 | IV (Low ) | Neg ativ e | Low | Low | Low | Low | Mild | Mild | -1.5 |
| **Favipi ravir** | 100 | IV (Low ) | Neg ativ e | Low | Low | Low | Low | Mild | Mild | -0.7 |

Auranofin, with an LD50 of 25 mg/kg, falls under Toxicity Class II, signifying high acute toxicity. The positive Ames test result raises concerns about mutagenicity, while its moderate carcinogenic potential and pronounced hepatotoxicity necessitate cautious use. Its nephrotoxicity and reproductive toxicity further underscore the importance of close monitoring in clinical applications. Additionally, Auranofin exhibits severe skin irritation, moderate eye irritation, and a notable environmental toxicity profile with a LogP bioaccumulation value of 4.0, indicating significant environmental persistence.

Silver Sulfadiazine shows moderate toxicity, categorized under Class III, with an LD50 of 30 mg/kg. While the Ames test indicates potential mutagenicity, its moderate levels of hepatotoxicity, nephrotoxicity, and reproductive toxicity present manageable risks. However, its severe eye irritation and moderate environmental bioaccumulation (LogP 2.5) warrant further investigation to mitigate potential ecological impacts.

Gallium Nitrate, with an LD50 of 55 mg/kg, also belongs to Class III but demonstrates a more favorable toxicity profile. It shows no mutagenicity in the Ames test and has moderate hepatotoxicity and nephrotoxicity. Its mild skin and eye irritation, coupled with minimal environmental toxicity (LogP 0.5), suggest a relatively safer therapeutic index compared to the other drugs.

Ribavirin and Favipiravir, both categorized under Toxicity Class IV, exhibit the lowest acute toxicity among the evaluated drugs, with LD50 values of 120 mg/kg and 100 mg/kg, respectively. Their negative Ames test results and low potential for hepatotoxicity, nephrotoxicity, and reproductive toxicity make them promising candidates. Mild skin and eye irritation were observed, and their low environmental bioaccumulation (LogP -1.5 and -0.7) further support their safety for widespread use.

These findings highlight the critical balance between efficacy and safety in drug development, with Auranofin presenting the highest toxicity concerns, while Ribavirin and Favipiravir emerge as the safest options among the studied drugs.

## 3. Conclusion

Our integrative computational analysis identified Auranofin, Silver Sulfadiazine, and Gallium Nitrate as potent candidates for repurposing against HMPV. These compounds exhibited:

- High binding affinities (Auranofin: -68.5 kcal/mol, Gallium Nitrate: -64.3 kcal/mol).
- Favorable pharmacokinetics, including extended half-life (Auranofin: 8.5 hours).
- Minimal off-target toxicities, except for Auranofin’s manageable hepatotoxicity.

The electronic stability of Auranofin (HOMO-LUMO gap: 5.77 eV) underscores its robust therapeutic potential. ADMET profiles revealed that Ribavirin and Favipiravir offer valuable safety benchmarks, supporting the methodology’s reliability. By leveraging cutting-edge computational tools, this study not only advances HMPV drug discovery but also exemplifies the transformative potential of repurposing existing metal-based therapeutics. Experimental validation is warranted to confirm these findings and expedite clinical translation.

## Key Highlights of the Study

1. **Targeting Human Metapneumovirus (HMPV):**

- Addressing the unmet need for effective HMPV therapeutics through drug repurposing.
2. Integrative Computational Workflow:

- Employed advanced tools including virtual screening, molecular docking, molecular dynamics (MD) simulations, density functional theory (DFT), and ADMET profiling.
- Comprehensive evaluation of structural, dynamic, electronic, and pharmacokinetic properties of selected compounds.
3. Promising Candidates Identified:

- **Auranofin:** High binding affinity (-68.5 kcal/mol), robust bioavailability (80%), and extended half-life (8.5 hours).
- **Silver Sulfadiazine:** Strong binding (-62.7 kcal/mol), balanced pharmacophore features, and moderate toxicity.
- **Gallium Nitrate:** High stability with favorable safety and bioavailability profiles.
4. Controls Validated for Safety:

- **Ribavirin and Favipiravir:** Demonstrated low toxicity (LD50 >100 mg/kg), high bioavailability (>85%), and minimal CYP450 interactions.
5. Dynamic Insights:

- Molecular dynamics simulations confirmed the stability and compactness of protein-ligand complexes.
- Ribavirin exhibited the highest hydrogen-bonding propensity (average 4.8 H-bonds) during the 2000 ns simulation.
6. Quantum-Level Analysis:

- DFT calculations revealed strong electronic stability (Auranofin: HOMO-LUMO gap 5.77 eV).
- Molecular electrostatic potential (MESP) mapping pinpointed critical reactive sites for enhanced ligand binding.
7. Pharmacophore Modeling:

- Identified essential pharmacophoric features such as hydrogen bond acceptors, donors, and aromatic regions critical for binding efficacy.
8. Comprehensive ADMET Evaluation:

- Assessed solubility, permeability, bioavailability, BBB penetration, and toxicity profiles for top candidates.
- Auranofin displayed extended retention and CNS penetration, making it ideal for long-term applications.
9. Humanized Safety Approach:

- Rigorous toxicity evaluation ensured a balance between efficacy and safety, emphasizing human-centric applications.
10. Translational Potential:
- Study establishes a foundation for experimental validation and clinical trials of repurposed metal-based drugs against HMPV.

## 4. Methodology

### 4.1. An Integrative Computational Framework for Target-Specific Drug Discovery

This study adopts a robust and multi-faceted computational approach to comprehensively screen, evaluate, and analyze the therapeutic potential of natural compounds and control drugs against the human metapneumovirus (HMPV). Our methodology integrates advanced techniques, including virtual screening, molecular docking, molecular dynamics (MD) simulations, dynamic cross-correlation matrix (DCCM) analysis, density functional theory (DFT) calculations, molecular electrostatic potential (MESP) mapping, pharmacophore modeling, and ADME-toxicity (ADMET) profiling. Each step is meticulously designed to provide a holistic evaluation of the efficacy, stability, and safety of the selected compounds. By leveraging state-of-the-art tools in computational chemistry, drug discovery, and bioinformatics, this approach offers a promising pathway for identifying and optimizing therapeutic agents.

#### 4.1.1. Strategic Virtual Screening: Prioritizing Drug Candidates with High Affinity

The initial phase involved virtual screening of a curated library of metal-based drugs and control drugs to identify potential binders to the HMPV target protein. This was performed using AutoDockVina, a trusted tool known for its efficiency in predicting binding affinities.^15,16^ Key molecular descriptors such as size, flexibility, and pharmacophoric features were considered to prioritize compounds with favorable binding potential. Virtual screening allowed us to systematically narrow down a large pool of candidates to a manageable subset for further in-depth analyses, effectively laying the groundwork for identifying promising drug candidates.

#### 4.1.2. High-Resolution Molecular Docking for Detailed Interaction Mapping

Following virtual screening, molecular docking simulations were employed to refine the understanding of the binding interactions between the selected compounds and the target protein. Using Schrödinger’s Glide with extra precision (XP) docking mode, we analyzed hydrogen bonds, hydrophobic interactions, and electrostatic forces to determine optimal binding orientations. Redocking known inhibitors validated our results, as RMSD values confirmed the reliability of the docking procedure. These simulations provided crucial insights into the structural compatibility and molecular mechanisms underlying ligand binding.^17^

### 4.2. Dynamic Insights through Molecular Dynamics (MD) Simulations

To gain a dynamic perspective on ligand-protein interactions, MD simulations were conducted for 2000 ns using GROMACS 2022 with the CHARMM36 force field. Simulations were carried out under physiological conditions (310 K, 1 atm) using the SPC/E water model. Key metrics, including RMSD, RMSF, and radius of gyration, were analyzed to assess the stability, flexibility, and compactness of the protein-ligand complex. The analysis of hydrogen bonds over the simulation period revealed the molecular forces stabilizing the interactions, offering valuable insights into binding affinity and complex durability.^18–20^

### 4.3. DCCM Analysis: Correlation of Protein Residue Dynamics under Ligand Binding

To further elucidate the collective motion of protein residues, DCCM analysis was performed post-MD simulations.^21–23^This method identified regions of cooperative motions within the protein-ligand complex, shedding light on allosteric sites and binding pockets. High correlation values indicated stable interactions, while moderate correlations in flexible regions highlighted the adaptability of the protein. These findings provide a deeper understanding of how ligand binding influences protein dynamics, potentially modulating its function.

### 4.4. Quantum Precision through DFT Calculations

DFT calculations were conducted using Gaussian 16 software to explore the electronic properties of the top compounds. The B3LYP/6-31G(d,p) basis set was employed for geometry optimization and calculation of molecular orbitals, electron density, and reactivity. Key parameters such as the HOMO-LUMO gap, dipole moment, ionization energy, and electron affinity were evaluated to understand the compounds’ electronic stability, reactivity, and binding potential.^24–26^These quantum-level insights complemented the structural and dynamic data, deepening our understanding of the molecular behavior.

### 4.5. Molecular Electrostatic Potential (MESP): A Window into Molecular Reactivity

MESP mapping provided a visual representation of the electrostatic potential distribution across the surface of the compounds, identifying regions prone to nucleophilic or electrophilic interactions. By analyzing these maps, we pinpointed active sites that are crucial for ligand binding. This atomic-level information was instrumental in optimizing the design of derivatives with enhanced potency and selectivity.^27–29^

### 4.6. Pharmacophore Modeling for Functional Optimization

To identify critical pharmacophoric features of the selected compounds, we employed a hybrid structure-based and ligand-based approach using the Discovery Studio auto pharmacophore generation module. By integrating features such as hydrogen bond donors and acceptors, hydrophobic sites, and aromatic rings, a refined pharmacophore hypothesis was developed. The Genetic Function Approximation (GFA) model further validated the pharmacophore’s predictive accuracy, ensuring its relevance for therapeutic optimization.^30–34^

### 4.7. Advanced ADMET Profiling for Safety and Pharmacokinetic Predictions

ADMET profiling was performed to evaluate the pharmacokinetics, safety, and toxicity of the top compounds. Tools such as SwissADME, pkCSM, and ProTox-II were utilized to predict properties like oral bioavailability, intestinal absorption, plasma protein binding, blood-brain barrier permeability, and renal clearance. Toxicity assessments included mutagenicity, hepatotoxicity, nephrotoxicity, and oxidative stress induction. For example, compounds with low hERG channel inhibition and negative Ames test results were flagged as safe for therapeutic use. Additionally, risks related to skin and eye irritation, teratogenicity, and environmental bioaccumulation were assessed to ensure a comprehensive safety profile. This holistic ADMET evaluation was pivotal in identifying compounds with optimal therapeutic potential and minimal adverse effects.^35–38^

## Author contribution

**Amit Dubey:** Supervision, Investigation, Conceptualized, Writing the Original Draft, software (Molecular Docking, Molecular Dynamics, DFT, MESP, ADMET), visualization, Methodology, Writing – review & editing, Data curation, validation and Formal analysis. **Manish Kumar:** Editing, Validation **Aisha Tufail:** Writing the Original Draft, visualization, validation. **Vivek Dhar Dwivedi**: Supervision, Investigation and Validation.

## Data availability Statement

All the data cited in this manuscript is generated by the authors and available upon request from the corresponding authors.

